# Multiple Pleistocene refugia for Arctic White Heather (*Cassiope tetragona*) supported by population genomics analyses of contemporary and Little-Ice-Age samples

**DOI:** 10.1101/2023.07.05.547859

**Authors:** Cassandra Elphinstone, Fernando Hernandez, Marco Todesco, Jean-Sébastien Légaré, Winnie Cheung, Paul C. Sokoloff, Annika Hofgaard, Casper T. Christiansen, Esther R. Frei, Esther Lévesque, Gergana N. Daskalova, Haydn J. D. Thomas, Isla H. Myers-Smith, Jacob A. Harris, Jeffery M. Saarela, Jeremy L. May, Joachim Obst, Julia Boike, Karin Clark, Katie MacIntosh, Katlyn R. Betway-May, Mats P. Björkman, Michael L. Moody, Niels Martin Schmidt, Per Molgaard, Robert G. Björk, Robert D. Hollister, Roger D. Bull, Sofie Agger, Vincent Maire, Liam Case, Greg H.R. Henry, Loren H. Rieseberg

## Abstract

**Aim:** Arctic plants survived the Pleistocene glaciations in unglaciated refugia, but the number of these refugia is often unclear. We use high-resolution genomic data from present-day and Little-Ice-Age populations of Arctic White Heather (*Cassiope tetragona*) to re-evaluate the biogeography of this species and determine whether it had multiple independent refugia or a single refugium in Beringia.

**Location:** Circumpolar Arctic and Coastal British Columbia (BC) alpine

**Taxon:** *Cassiope tetragona* L., subspecies *saximontana* and *tetragona,* outgroup *C. mertensiana* (Ericaceae)

**Methods:** We built genotyping-by-sequencing (GBS) libraries using *Cassiope tetragona* tissue from 36 Arctic locations, including two ∼250-500-year-old populations collected under glacial ice on Ellesmere Island, Canada. We assembled a *de novo* GBS reference and called variants in dDocent. Population structure, genetic diversity, and demography were inferred from PCA, ADMIXTURE, fastsimcoal2, SplitsTree, and several population genomics statistics.

**Results:** Population structure analyses identified 4-5 clusters that align with geographic locations. Nucleotide diversity was highest in Beringia and decreased eastwards across Canada. Demographic coalescent analysis of the site-frequency-spectrum dated the following splits from Alaska: BC subspecies *saximontana* (6 mya), Russia (1.5 mya), Europe (>300-600 kya), Greenland (100 kya). Northern Canada populations appear to be from the current interglacial (7-9 kya). Genetic variants from Alaska appeared more frequently in present-day than historic plants on Ellesmere Island.

**Conclusions:** Demographic analyses show BC, Alaska, Russia, Europe, and Greenland all had separate refugia during the last major glaciations. Northern Canadian populations appear to be founded during the current interglacial with genetic contributions from Alaska, Europe, and Greenland. On Ellesmere Island, there is evidence for continued, recent gene flow with foreign variants introduced in the last 250-500 years. These results suggest that a re-analysis of other Arctic species with shallow population structure using higher resolution genomic markers and demographic analyses may help reveal deeper structure and other circumpolar glacial refugia.

## Introduction

The Arctic flora has a complex biogeographic history due to repeated glacial cycles causing extreme range expansions and contractions throughout the Pleistocene (Hultén, 1937; Dyke & Prest, 1987; Abbott & Brochmann, 2003; Hughes et al., 2016; Dalton et al., 2020). During each glacial period, Arctic species survived in isolated non-glaciated refugia where they had time to diverge from each other. Following this isolation, range expansions during post-glacial periods led to admixture among populations (Hultén, 1937). While complex biogeographic histories can be difficult to deduce from fossil data, genomic data offer a means to infer refugia, their relative ages, potential colonization routes, and to estimate gene flow (Wang et al., 2021). Genomics studies have huge potential in rapidly changing Arctic ecosystems to help determine complex past population demography and predict future adaptation (Wullschleger et al., 2015). Few studies have tested the ages of potential Arctic plant refugia using high resolution genomic data (Ikeda et al., 2017) and there is still much uncertainty as to where and when these refugia formed for various species. By combining broad geographical sampling, high resolution genomic markers, and historical samples, we can begin to infer the age of Arctic refugia, their geographic locations, and timing of recent gene flow in Arctic plant populations (Wang et al., 2021).

For nearly a century, Beringia (the region between eastern Siberia and western Yukon) has been proposed an unglaciated refuge and the main origin of Arctic plant species dating back as far as 5.3 million years ago (mya) (Abbott et al., 2000; Brochmann & Brysting, 2008; Hultén, 1937). Genetic diversity there is generally higher compared to the rest of the Arctic, suggesting Beringia lineages are older and have had more time to diversify, but details for individual species remain uncertain (Abbott & Brochmann, 2003; Eidesen et al., 2013). However, there is also evidence for other Pleistocene-aged refugia throughout the Arctic. Other possible refugia for Arctic plants during the last major glaciation were mountains in northwestern USA, Western Europe, and South-Central Asia (Pellissier et al., 2016). Nunataks, unglaciated rocky outcrops in ice fields, have also been proposed as potential cryptic refugia where Arctic/alpine species may have survived in small, isolated populations during the glaciations (Beatty & Provan, 2010; Westergaard et al., 2011). Fossils and pollen records support that many High Arctic plant species had circumpolar distributions at various points in time since the polar tundra biome initially formed about 2-3 mya (Brochmann & Brysting, 2008). However, most previous molecular datasets on Arctic plants used highly conserved chloroplast DNA sequences or a small number of AFLPs, so fine-scale resolution of genomic clusters and estimation of demographic history was not possible. One exception is a study in *Kalmia procumbens* that estimated divergence times to be during the last interglacial from Beringia using RADseq data (Ikeda et al., 2017).

Since 1950-1960, following the Little Ice Age (ca. 1450-1850 CE), the polar-frozen-based glaciers in the High Arctic have been retreating. As the ice retreats, historic plants are uncovered essentially intact (Bergsma et al., 1984; Jones & Henry, 2003). At Alexandra Fiord, Ellesmere Island, in the last 60 years the ice has retreated approximately 300 m from its maximum extent (O’Kane, 2018). *Cassiope tetragona* is one of the best-preserved species to emerge from under the ice, and thus we have used these historic populations to extend our genetic sampling backwards in time.

*Cassiope tetragona* (L.) D. Don is a diploid (2n=26), perennial, ecologically well-studied (Havstrom et al., 1995; Mallik et al., 2011; Molau, 1997; Rayback & Henry, 2006) Arctic/alpine species, known as Arctic White Heather in English, and *qijuktaat* meaning “for fire building” in Inuktitut (Aiken et al., 2007). Two subspecies of *C. tetragona* have been described: ssp. *tetragona*, which has a circumpolar distribution; and ssp. *saximontana*, which is found in Western Canada. *Cassiope tetragona*is insect pollinated (Alsos et al., 2013) but may also self. Small seeds enable wind dispersal (Eidesen et al., 2007), facilitating widespread colonization. Under experimental conditions, *C. tetragona* has low seed germination (7.8%) (Alsos et al., 2013), possibly hampering post-dispersal establishment. *Cassiope tetragona* does not currently have a reference genome. Previous RADseq data on the *Cassiope* genus showed evidence that the genus originated in the north (Siberia) and later spread southwards (through the Himalayan-Hengduan Mountains) (Hou et al., 2016). *Cassiope tetragona* is believed to have split from its sister species *C. mertensiana* (Bong.) G. Don for about 18.6 mya (CI: 14.8 - 37.2 mya; Kumar et al., 2017).

Previous phylogenetic analyses of the Arctic populations of *C. tetragona* showed shallow structure, suggesting that the species may have expanded its range eastward out of a Beringian refuge across to Europe as recently as the last interglacial period (<11 thousand years ago (kya)) (Eidesen et al., 2007). They also suggested a westward expansion to Siberia in the mid-to-late Pleistocene. However, due to the use of relatively limited genotypic information (265 AFLP markers in 56 populations) that may have obscured more complex patterns of genetic diversity, the presence of additional refugia could not be ruled out.

Here, we re-investigate the biogeography of *C. tetragona* from 36 extant pan-Arctic populations, two historic (250-500 year-old) populations sampled from under glacial ice in the Canadian High Arctic and an outgroup (*C. mertensiana*) using up to 26,350 single nucleotide polymorphisms (SNPs). The main goal of our study was to determine if the current circumpolar population of *C. tetragona* derived from recent expansion out of a Beringian refuge, or if *C. tetragona* populations could have persisted in several other glacial refugia during the Pleistocene glaciations. Additionally, we also investigate if and how Pre-Little-Ice-Age populations differ from present-day populations.

## Materials and Methods

### Sample collection, DNA extraction

During summer of 2017, we, researchers primarily associated with the International Tundra Experiment (ITEX), a collaborative international group of researchers studying tundra ecosystems, (Henry et al. 2022), collected *C. tetragona* leaf tissue on silica from 36 pan-Arctic and alpine locations (Table S3, Figure 1B). Two historic populations (Alexandra Fiord and Sverdrup Pass, Ellesmere Island, Nunavut, Canada) were retrieved from retreating glacial ice. It was observed that the glacier coverage came quickly (possibly in the middle of summer - since many of the plants we found frozen were in flower (not having set seeds yet). Two populations of *C. mertensiana* from the alpine in southwestern British Columbia (BC) served as an outgroup (MER). All samples were collected as described in Appendix S1-Field Sampling in Supplementary Materials. Samples were collected from all countries in *C. tetragona*’s range. Population codes were assigned according to the geographic location of the sampling site (Table S3).

**Figure 1:**
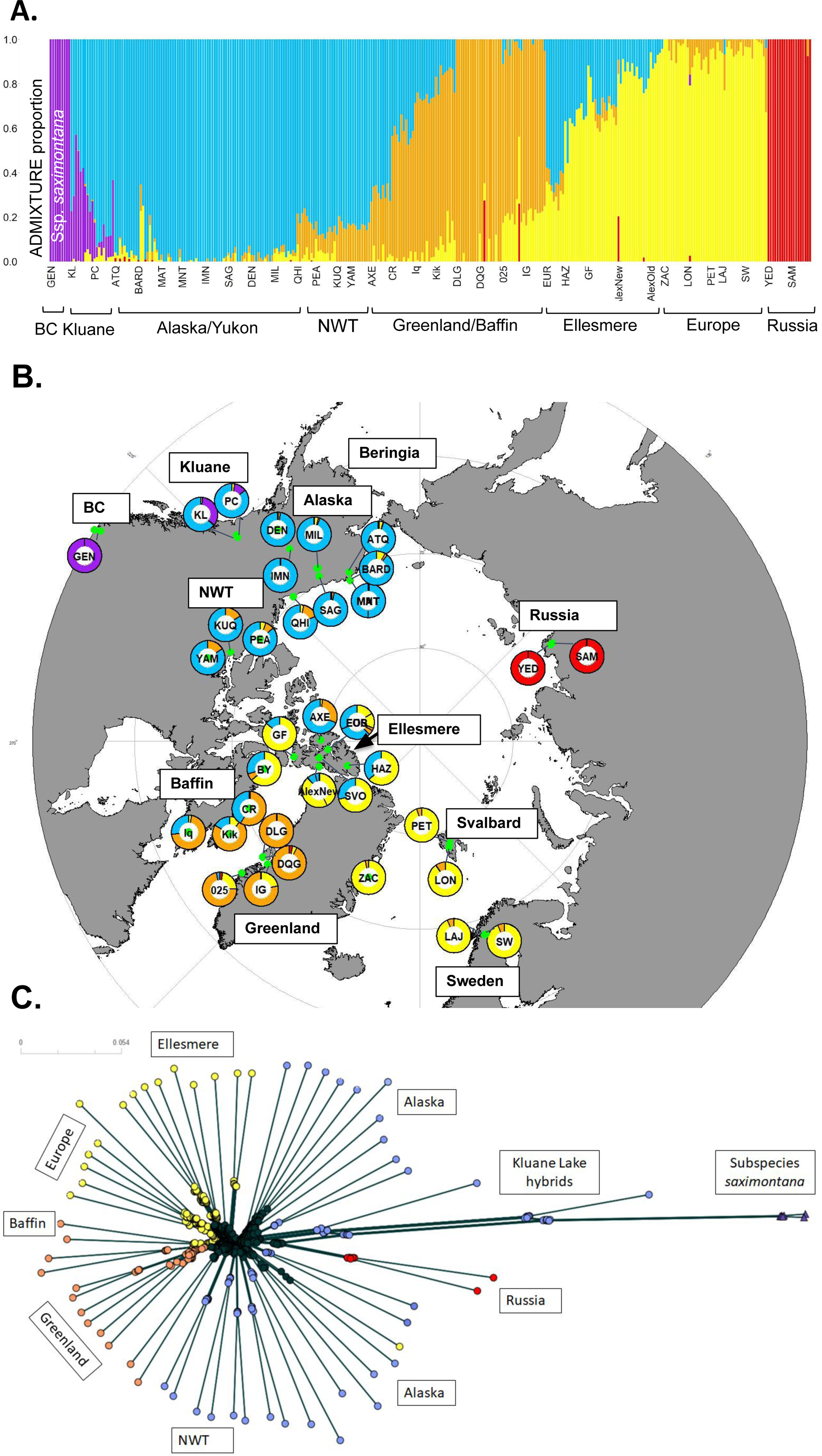
Population structure. Relationship between populations using ADMIXTURE results based on 9285 SNPs and 330 individuals. **(A)** Barplot shows individual genotypes as vertical bars ordered by populations and grouped by genetic cluster. Colours represent the proportion of that individual’s genotype associated with each cluster (Top: K=5, CV error=0.262). **(B)** Geographic map of collection sites. Each site is represented by a pie chart that is colour-coded by its ADMIXTURE groups (K=5). See Table S3 for information about all sample locations. **(C)** SplitsTree neighbor-net tree network built with the uncorrected P distance using two individuals randomly selected from every population (see Table S3 for collection location of each 3-letter code). Colours based on results from K=5 ADMIXTURE groups. Network based on 25 572 SNPs and 60 individuals. Scale bar shows the uncorrected P distance.

DNA was extracted from 20 mg of dried leaf tissue from each of the 5 to 12 individuals per site using a 3% CTAB protocol (Zeng et al., 2002) that was modified for high phenol content and acidity in *C. tetragona* leaves (Appendix S1-Modified 3% CTAB). The modified CTAB protocol was also used to extract DNA from 24 historic plants. In all, DNA was obtained from 387 samples, which were subsequently prepared for sequencing.

### Sequencing libraries

Genotyping by sequencing (GBS) was selected as a cost-effective way to obtain enough SNPs to differentiate populations in recently geographically expanded species such as *C. tetragona*. GBS was conducted using New England Biolabs PstI Hi-Fidelity and New England Biolabs MspI restriction enzymes following a modified version of the (Elshire et al., 2011) protocol (Appendix S1-Genotyping by sequencing library preparation). After a polymerase chain reaction (PCR) enrichment step, pooling, size selection of 300-500 base pair (bp) fragments, and enzymatic depletion of repetitive sequences (Moyers et al., 2017), enriched libraries for 371 samples were successfully prepared, including 10 of 12 historic samples from Alexandra Fiord and all 12 samples from Sverdrup Pass (16 of the 387 samples failed to amplify). Libraries were sequenced at the Centre d’Expertise et de Services Génome Québec (CESGQ) on two lanes of the Illumina HiSeq4000, generating 571 million 150 bp paired end reads in total. Sequence data are available in the GenBank/SRA database under accession number SUB11222726 and BioProject ID PRJNA824830.

### Data processing

All computations were run on the WestGrid Compute Canada clusters. Detailed data processing notes and scripts can be found here: https://github.com/celphin/Population_genomics_Cassiope. First, fastq files were de-multiplexed using a Perl script from Owens et al. (2016) and then loaded into dDocent 2.9.4 (Puritz et al., 2014). The ends of reads with a quality score below 20 or an average quality score that was <10 in a sliding window of 5 bp were trimmed with Trimmomatic (Bolger et al., 2014).

Since a reference assembly was not available for *C. tetragona*, we used GBS reads from a total of 55 *C. tetragona* individuals (from ATQ, BARD, LAJ, PET, SAM, YED; see Table S3) to build a *de novo* GBS assembly for paired end reads using the program CD-HIT (LaCava et al., 2020) and Rainbow (Chong et al., 2012) through the dDocent 2.9.4 assembly pipeline. A c-parameter (% similarity to cluster) of 0.95 was used to cluster reads. To be included in the assembly, reads had to have a within individual coverage of 3 and a between individual coverage of 5. Historic plants were excluded from the assembly to avoid the inclusion of contaminant bacterial DNA fragments in the assembly (see below). Similarly, *C. mertensiana* individuals were excluded from the assembly to prevent the generation of duplicated contigs for divergent regions. We then mapped reads from all *C. tetragona* individuals to our *de novo* GBS assembly using BWA-MEM (Li & Durbin, 2009) through the dDocent pipeline. Total mapping scores were assigned to each read based on a match score of 1, a mismatch score of 4 and a gap penalty of 6 (the conservative default values for BWA mem). Our outgroup samples from BC, *C. mertensiana* and ssp. *saximontana* were mapped with slightly more relaxed parameters of match score of 1, a mismatch score of 3 and a gap penalty of 5. Finally, variant calling was done on all individuals via FreeBayes (Garrison & Marth, 2012) within the dDocent pipeline (using the dDocent 2.9.4 default parameters and included both SNPs and insertions/deletions). High levels of DNA degradation in historic and ancient DNA samples can often lead to high levels of bacterial contamination in sequencing libraries. Bacterial vs eukaryotic reads were identified by blasting each read to the NCBI database and calculating the percentage of total reads that mapped to Bacteria vs Eukaryotes. The original percentage of bacterial reads in the historic samples can be found in Table S5; such reads were removed when mapping GBS data from the historic samples to the *de novo* reference genome.

### SNP Filtering

We filtered all variants in the raw variant call file (vcf) using vcftools 0.1.16 and plink 1.9b_6.21-x86_64 (Danecek et al., 2011). First, all variants that were heterozygous in >60% of individuals (using plink - -hardy option) across all populations were removed since these were likely mapping errors resulting from a *de novo* reference genome that may not have accurately resolved duplicated regions of the genome. Next, using vcftools, all variants were filtered based on the following criteria in order: quality score below 30, indels, non-biallelic SNPs, called in less than 90% of individuals, and with a minor allele frequency of <1%.

Filtering for linkage disequilibrium (LD) and individuals with missing data varied. Linked SNPs were removed (vcftools -- thin 300 (length of the longest contig)) for all analyses except the population statistics, SplitsTree and ABBA BABA test. We also removed individuals missing more than 10% of the variants found across all sites and the outgroup individuals (with poor mapping) from all analyses except the demographic modeling where we projected the data to 10 individuals per group (for the demography we retained the outgroup and any individuals missing up to 30% of SNPs).

### Population structure

To visualize variation and clustering among individuals and sites, principal component analysis (PCA) was performed in R 4.2.2 using the packages SNPRelate (Zheng et al., 2020) and tidyverse (Wickham et al., 2019).

ADMIXTURE 1.3.0 (Alexander & Lange, 2011) was used to assign potential genome ancestry to SNPs using a model-based approach with the assumption that ancestral populations were in Hardy-Weinberg Equilibrium (HWE). The termination criterion was set to when the log likelihood changes between iterations fell below 10^-10^. The tenfold cross validation (CV) error was recorded for each value of K (1 to 14) tested. The last large drop in CV error was used to select the K value best representative of the number of clusters in the data. All pairwise F_ST_ values between the identified clusters were calculated.

### Population statistics

Vcftools 0.1.16 was used to calculate general population statistics (expected and observed homozygosity, inbreeding coefficient (F_IS_), nucleotide diversity (π), F_ST_, and Tajima’s D). Each sampling location was grouped together as a “population”. Nucleotide diversity (π) was calculated for each variant site using the ‒site-π option and across 150 bp windows using the ‒window-π option in vcftools for each location. These π values were summed to obtain a single sites-π and windows-π summed measure for each location. Descriptions of other population statistics (expected and observed homozygosity, Tajima’s D, and the inbreeding coefficient (F_IS_)) can be found in Appendix S2 - Population statistics. For nucleotide diversity and other population statistics, maps of the average values per location were made using the PlotSvalbard package (Vihtakari, 2020).

If populations began diversifying at each location beginning when it was ice-free, time since the most recent deglaciation should be associated with genetic diversity measures. To investigate this, values of π for the Arctic populations (ssp. *tetragona*) were compared with dates at which the last glacial maximum ice sheets retreated at each location. Deglaciation dates were obtained from a polygon shapefile in (Dalton et al., 2020) using QGIS 2.18.10 (http://qgis.org) for North America; for the European sampling sites, approximate dates were selected from figures in (Hughes et al., 2016).

F_ST_ for all pairs of sampled populations were calculated in vcftools with the --weir-fst-pop option. F_ST_ values comparing the genetic distances between all sampling locations were used to investigate isolation-by-distance relationships. A Mantel test, limited to the *C. tetragona* ssp. *tetragona* populations, was conducted (mantel.rtest in ade4 R package (Thioulouse et al., 2018), R 3.5.0) with 9999 replicates.

### Neighbor-net tree and maximum likelihood tree

A neighbor-net tree was built with SplitsTree5 (Huson, 1998; Huson & Bryant, 2006). We used PGDSpider 2.1.1.5 (Lischer & Excoffier, 2012) to convert our vcf file into a standard binary nexus file. For this, both heterozygotes and homozygotes for the novel allele were labeled 1, while individuals homozygous for the reference allele were labeled 0. In SplitsTree5, uncorrected P-distances were calculated and splits were determined using Neighbor Net and Splits Network Algorithm for two randomly selected individuals from each sampling location listed in Table S3.

### Demographic models (site frequency spectrum)

To estimate the relative ages, migration, and population sizes of the various admixture clusters we ran a coalescent demographic analysis on the folded site frequency spectra using fastsimcoal2 (Excoffier et al., 2021). We tested four different models each including 5 clusters: ModelA: *C. mertensiana*, *C. tetragona* ssp. *saximontana*, Russia, Alaska, and Europe; ModelB: *C. mertensiana*, Russia, Alaska, Europe and Greenland; ModelC: Russia, Alaska, Europe, NWT and Nunavut; and ModelD: Alaska, Europe, Greenland, NWT and Nunavut (Figure S5). To estimate parameters, we set fixed known divergence time estimates. In the first two models, we used the split between *C. tetragona* and *C. mertensiana* (18.6 mya) (Kumar et al., 2017), while in the third and fourth models, we assumed that populations in Northern Canada (NWT and Nunavut) were formed since the last major glaciation (7-9 kya, respectively). Divergence times were expressed in generations; *Cassiope tetragona* was assumed to have a generation time of 10 years. Its generation time is not known but observationally it does not flower for at least the first five years of its life and can live to be up to 70-150 years old (Rayback & Henry, 2006; Weijers et al., 2017). For the first two models, we selected individuals with >95% of membership to their group in ADMIXTURE, excluding hybrid individuals. Each group was projected to ten individuals (--proj 20 in easySFS) (Gutenkunst et al., 2009), with the exception of *C. mertensiana*, which only had 7 non-hybrid individuals. Model3 and Model4 included more recently formed Canadian hybrid populations, so to account for the admixed ancestry of these populations, we included in the models a bidirectional gene flow matrix. Each model was run 50 times. Within each run 100 000 simulations were done, each with 48 optimization iterations. The best run of each model was selected based on the maximum estimated likelihood. Models were compared with each other by checking the difference between the maximum estimated likelihood and the model maximum observed likelihood. Site frequency spectra were plotted for the raw data in python 3.10 with dadi (Gutenkunst et al., 2009).

### Historic samples

To determine the ages of the historic tissue from under the glacial ice, leaf/stem tissue from five of the historic samples were C-14 dated using Accelerator Mass Spectrometry (AMS) at the André E. Lalonde AMS Laboratory, University of Ottawa.

To test for recent admixture between two major genetic clusters, present-day and historic populations at Alexandra Fiord were used. We employed ABBA-BABA tests in Dsuite (Figure 5C) (Durand et al., 2011; Green et al., 2010; Malinsky et al., 2021) across four populations: P1, P2, P3, and the outgroup (O), related by the phylogeny (((P1, P2), P3), O). SNPs for this analysis were filtered to have as many variants as possible to compare between individuals from the outgroup (non-hybridized individuals), AlexOld and AlexNew samples. SNPs were filtered out if they: had a quality score <30, were indels or non-biallelic SNPs, and called in less than 95% of individuals (from the outgroup, AlexNew and AlexOld). The allele carried by the outgroup (seven samples of *C. mertensiana*) was designated as the ancestral allele (A) and the derived allele was designated as B. Under the null hypothesis, which assumes no gene flow after P1 and P2 split, the ABBA (B shared by P2 and P3) and BABA (B shared by P1 and P3) patterns are expected to occur with equal frequency due to incomplete lineage sorting. A significant increase in ABBA or BABA is consistent with introgression between P3 and either P1 (ABBA < BABA, negative D values) or P2 (ABBA > BABA, positive D values). In each scenario, P1 and P2 were represented by the historic (AlexOld) and present-day (AlexNew) samples, respectively, and as P3 we used multiple populations from Alaska (ATQ, BARD, DEN, IMN, MIL, MNT, and SAG) as well as an artificial population formed by 10 individuals from the Alaska genetic cluster (admixture membership > 0.99).

### Flow cytometry

Flow cytometry was run to estimate *C. tetragona*’s genome size and check the ploidy of a few populations on Disko Island, Greenland. We cut up silica dried *C. tetragona*and *Solanum lycopersicum* (genome size standard variety grown from seed) leaf tissue in our nuclei extraction buffer, washed and fixed the nuclei before dying with propidium iodide. Flow cytometry was run on a Cytoflex flow cytometer in the UBC Biomedical Research Center. Our detailed flow cytometry protocol can be found in Appendix S1-Flow cytometry.

## Results

### Filtering

Variant calling was performed on GBS data for 371 *Cassiope* samples from 36 populations (Table S3), resulting in a total of 559 815 variant sites. The 10 *C. mertensiana* individuals were only used as an outgroup for the ABBA BABA tests and in demographic modelling because read mapping to the *de novo C. tetragona* reference genome was poor for many of these individuals (missing data amounts are listed in Table S4).

### Population structure

In PCAs, approximately 27% of genetic variation in the dataset was associated with the first principal component (PC1) separating the subspecies *C. tetragona* ssp. *saximontana* (purple) from the rest of the Arctic *C. tetragona* ssp. *tetragona* (Figure 2A, Table S6). Individuals from Kluane Lake, Yukon (blue) stand out as apparent hybrids between the two subspecies. Approximately 6% of the variation occurs along PC2 separating Russia (red) from the rest of the Arctic. There appear to be a few individuals from Greenland (orange) found between Russia (red) and the rest of the Arctic. Despite the low amount of variation explained by PC3 (2.63%) and PC4 (1.63%), PC3 separates Alaska (blue) from Russia, Europe, and Greenland (red/yellow/orange) and PC4 splits Greenland (orange) and Europe (yellow) in opposite directions from Alaska and Russia.

**Figure 2:**
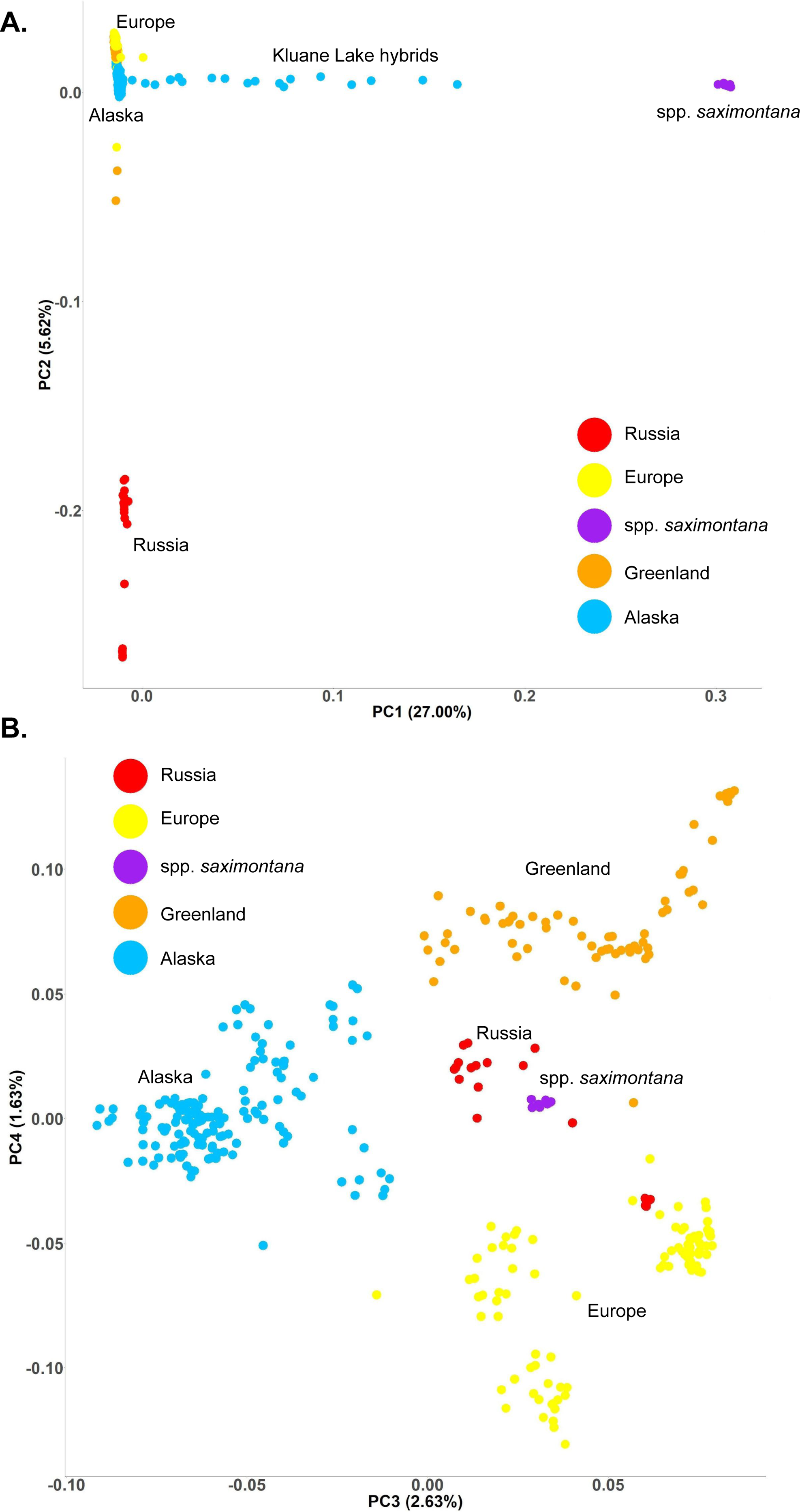
Principal component analysis. (PCA) run on 9 285 variants and 349 *Cassiope tetragona* individuals. Individuals are colour-coded according to the K=5 ADMIXTURE groupings (outgroup *C. mertensiana* is excluded). Percentage of variance accounted for by each component shown with the axis label. (A) The second principal component (PC2) versus PC1. (B) PC4 versus PC3.

ADMIXTURE clustering results for K=1-14 are largely consistent across multiple tests (with K=5 consistently showing the last large drop in CV scores; Table S7). At K=5, ADMIXTURE results suggest unique clusters currently in BC, Russia, Alaska, Europe, and Greenland. Populations in Northern Canada (NWT, Baffin and Ellesmere) appear to have mixed ancestry from Alaska and Europe (Figure 1a).

A neighbor-net tree built using SplitsTree5 (Figure 1C) with 25 572 SNPs and 60 individuals agrees with the findings from the PCA (Figure 2) and ADMIXTURE analysis (Figure 1A). A long branch separates the outgroup*, C. mertensiana* (MER), from the *C. tetragona*Arctic populations. Geographic location is associated with genealogical relatedness, with individuals from different regions forming their own distinct clades. Populations in the Kluane region in SW Yukon (KL and PC sites) gradually transition from the Arctic ssp. *tetragona* cluster to *C. tetragona* ssp. *saximontana* (GEN) populations in southern British Columbia. This pattern is consistent with the ADMIXTURE and PCA results and suggests admixture between ssp. *tetragona* and ssp. *saximontana* in the Kluane region.

### Population statistics (Nucleotide diversity (π), F_ST_ and genetic distance)

The windows and site-based approaches for calculating nucleotide diversity gave comparable results. The statistics were based on 26 350 SNPs after filtering. A significant positive relationship was found between ice retreat times and the amount of diversity both when including individuals with subspecies *saximontana*ancestry and when not (without *saximontana*, n=31, r^2^=0.31, p-value=0.00032, Figure S8). This relationship was driven by the higher diversity and older populations in Beringia because when samples from Beringia were removed the relationship was no longer significant (n=31, r^2^=0.072, p-value=0.095) (Figure 3A, Table S8).

**Figure 3:**
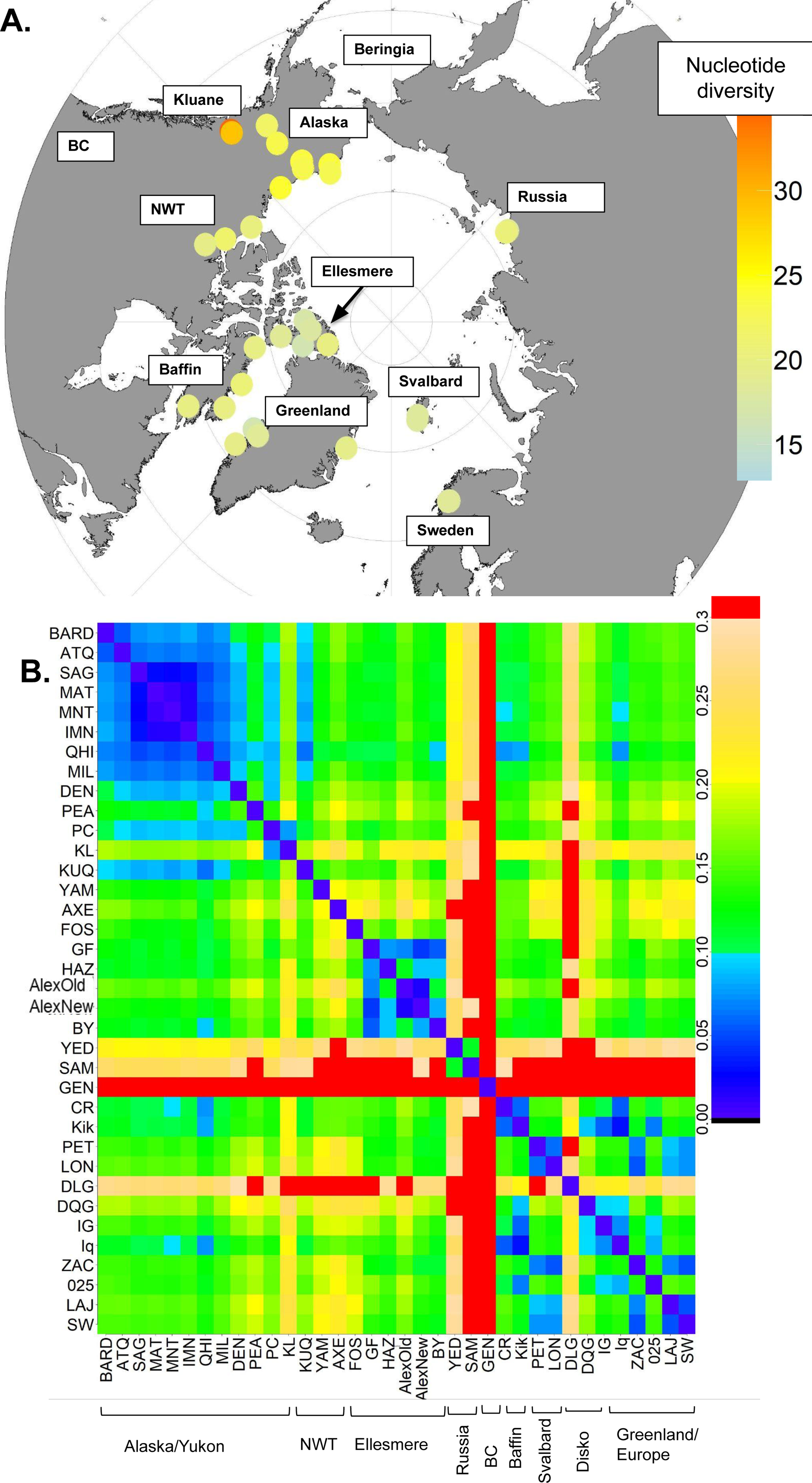
Population statistics. **(A)** Sum of window-based nucleotide diversity (π) mapped for each Arctic population (See Table S3 for names and location information of sample sites). **(B)** F_ST_ values for all pairs of sampled locations (ordered by distance from Barrow, Alaska). F_ST_ values >0.3 are shown in red (British Columbia *saximontana* population = GEN).

Generally, low F_ST_ values (near zero) indicate largely random mixing among populations, while values closer to one are indicative of isolated populations connected by little gene flow. F_ST_ was calculated for all the geographically separated sampling locations (Figure 3B, for all *C. tetragona* populations listed in Table S3) and for the regional ADMIXTURE clusters for K=5 (Figure S6A). As expected, the largest F_ST_ values were found when comparing the subspecies *C. tetragona* ssp. *tetragona* in the Arctic and the ssp. *saximontana*cluster in BC (GEN) (F_ST_∼0.6) (Figure 3B). Within spp. *tetragona*, the Russian populations (SAM and YED) showed the highest F_ST_ values (∼0.3) in pairwise comparisons followed by the Disko Island in Greenland (F_ST_∼0.2; DLG, Table S3, dark green). The rest of the Arctic clusters (Alaska, Yukon, NWT, Nunavut, Greenland, Svalbard and Sweden) all had lower F_ST_ values (F_ST_<0.15) when compared with geographically close populations than with populations further away. Inbreeding coefficients were also determined with a slightly negative mean but not significantly different from zero (Figure S7).

Genetic distance appears to increase with geographic distance (Mantel test, p-value=1.0x 10^-4^) (Figure S6B). The relationship between minimum F_ST_ and geographic distance appears positive, with close populations mostly having F_ST_ values near zero (except for Disko Island) and populations 5 000 km apart having a minimum F_ST_ value of 0.15.

### Demographic analysis (fastsimcoal2)

In all four models, genetic groups diverged well before the current interglacial (Table 1, Figure S9, S10, S11), supporting the presence of independent refugia in BC (known subspecies), Alaska, Russia, Europe, and Greenland. Divergence times were mostly consistent with genetic differentiation values between groups (F_ST_). For example, in our model *C. tetragona* ssp. *saximontana*diverged first from Alaska *C. tetragona*spp. *tetragona* about 6 mya (F_ST_=0.75). Russia diverged next approximately 1.5 mya (F_ST_=0.22), followed by Europe 200-600 kya (F_ST_=0.14), and then Greenland split from Europe 100 kya (F_ST_=0.20) (Figure 4, Table 1). The lower F_ST_ value between Europe and Alaska may be explained by the ongoing gene flow through the Canadian hybrids. Estimations of effective population sizes and divergence times between the four models were largely consistent within an order of magnitude (Table 1).

**Figure 4:**
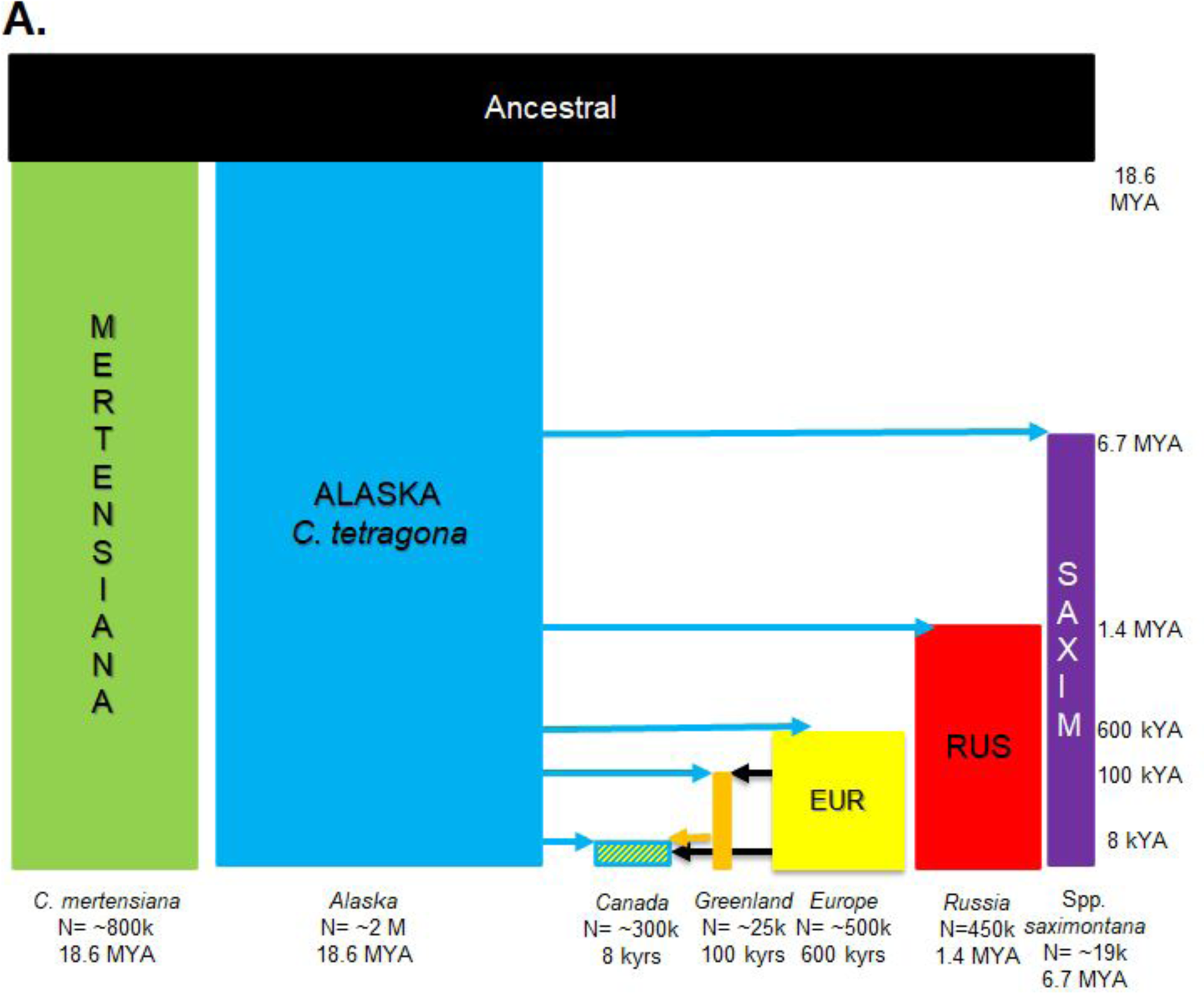
Coalescent demographic analysis results from fastsimcoal2. Combined schematic of all four models in Figure S5 and approximate results. See Table 1 for exact model output parameters. The size of the rectangles approximately represents the age of the populations (vertical axis) and the effect population size (horizontal axis). Old, large populations are larger than small, young populations. Each population is coloured to match its dominant K=5 ADMIXTURE group (matching Figure 1).

**Table 1:**
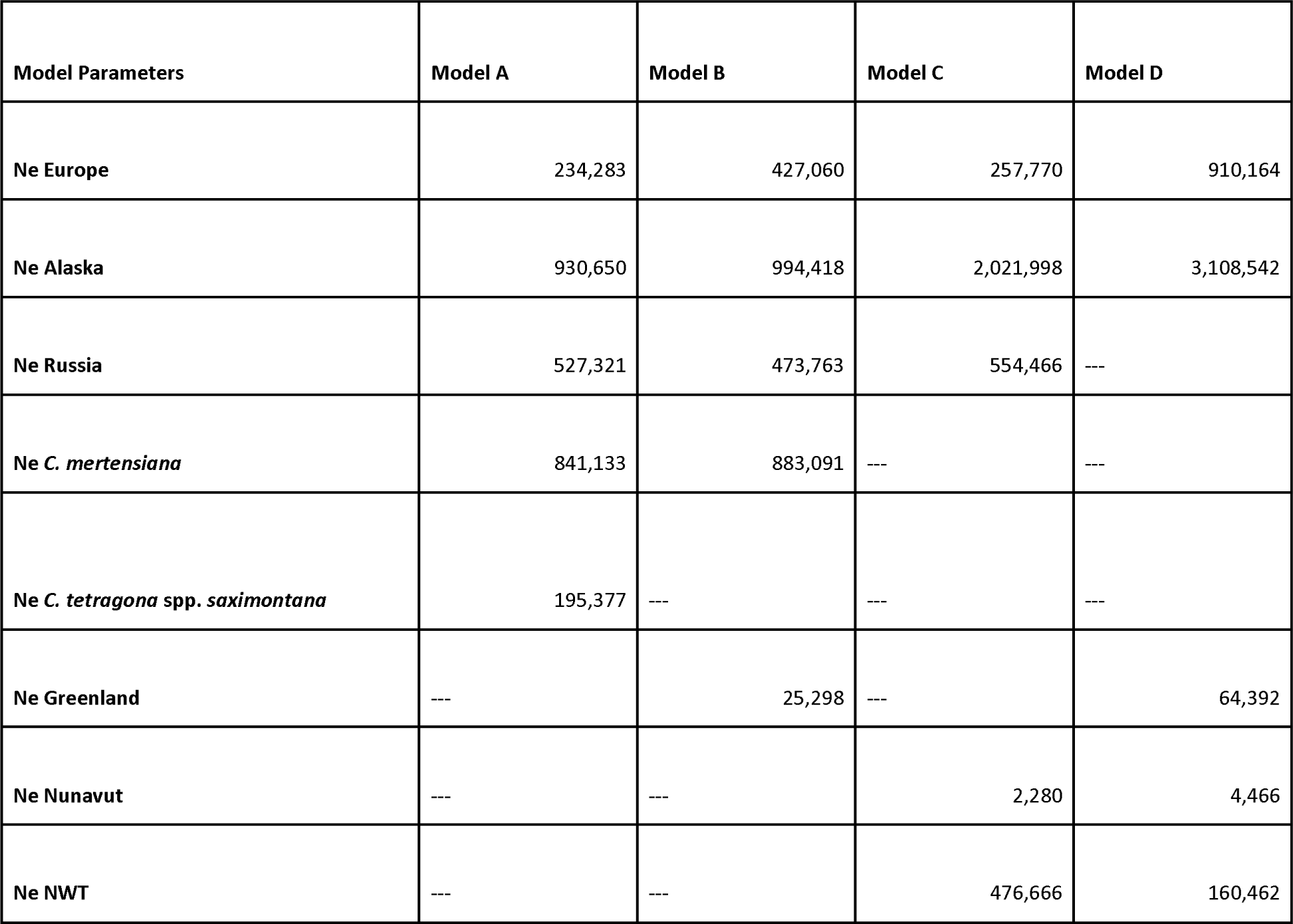
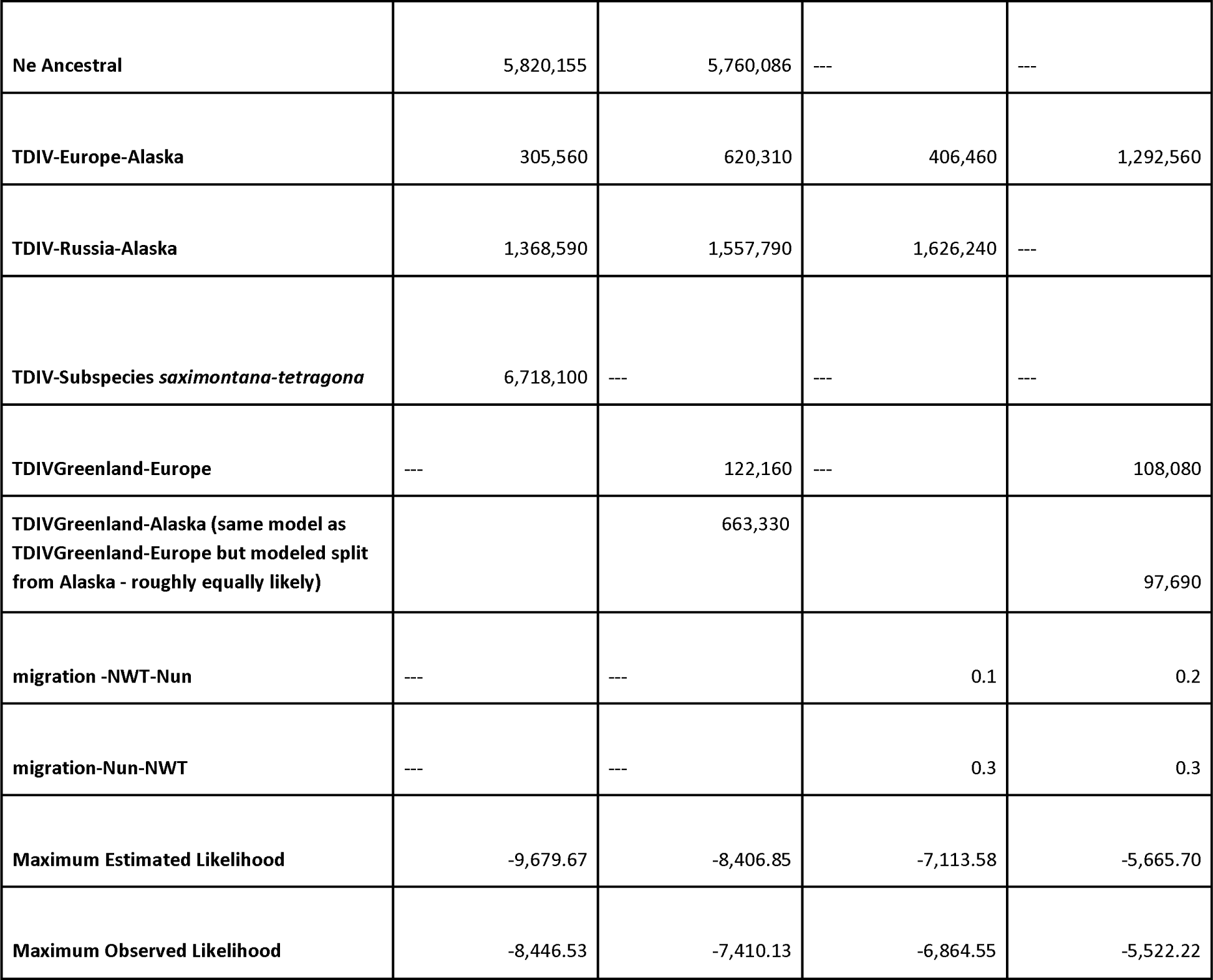
Coalescent demographic analyses using fastsimcoal2 results. Models A and B used a fixed split time of the sister species *Cassiope tetragona* and *C. mertensiana* at 18.6 mya while models C and D used the founding time of the Northwest Territories (NWT) and Nunavut to be 7000 and 9000 years ago, respectively. Ne is the estimated effective population size, TDIV is the time since divergence. All times are in years based on an estimated generation time of 10 years.

### Flow cytometry (genome size)

We determined the genome size of both populations found on Disko Island, Greenland (DLG and DQG) to test if a difference in ploidy was causing the high F_ST_ values between these very close populations. Flow cytometry results (Table S9) showed that *Cassiope tetragona* has a haploid genome size of roughly 1.6 Gbp and there were no clear genome size differences between the two populations on Disko Island.

### Historic samples

Carbon dating estimated samples from under the Twin Glaciers at Alexandra Fiord on Ellesmere Island (AlexOld) to be between 270 and 430 years old, and samples from Teardrop Glacier at Sverdrup Pass (SVO) to be 290-520 years old (exact values Table S10). Glaciers in North America began expanding in the mid-1500s (Forbes et al., 2020) and this expansion stopped in the late 19^th^ to early 20^th^ century.The historic samples therefore were likely living plants existing just prior to the glacial expansion. ADMIXTURE results comparing only the historic and present-day Alexandra Fiord populations (AlexNew and AlexOld) indicated that present-day samples have a higher proportion of ‘Alaskan’ variants compared to the historic populations (Figure 5A,B). A Mann Whitney U test confirmed this significant difference in ancestry proportions. Historic and present-day ancestry comparison for the Alaskan/Ellesmere cluster had a p-value = 0.0046 and the European/Ellesmere cluster had a p-value= 0.0023. ABBA-BABA test results consistently show that modern samples from Alexandra Fiord share more alleles with populations from Alaska than historical samples (Figure 5C). Tests were significant using 29 824 variants for various Alaskan populations selected as P3 (Z = 3.25; p-value= 0.001; fraction of introgression f_4_-ratio = 0.15; for P3=Alaska generally, other P3 population values can be found in Table S11).

**Figure 5:**
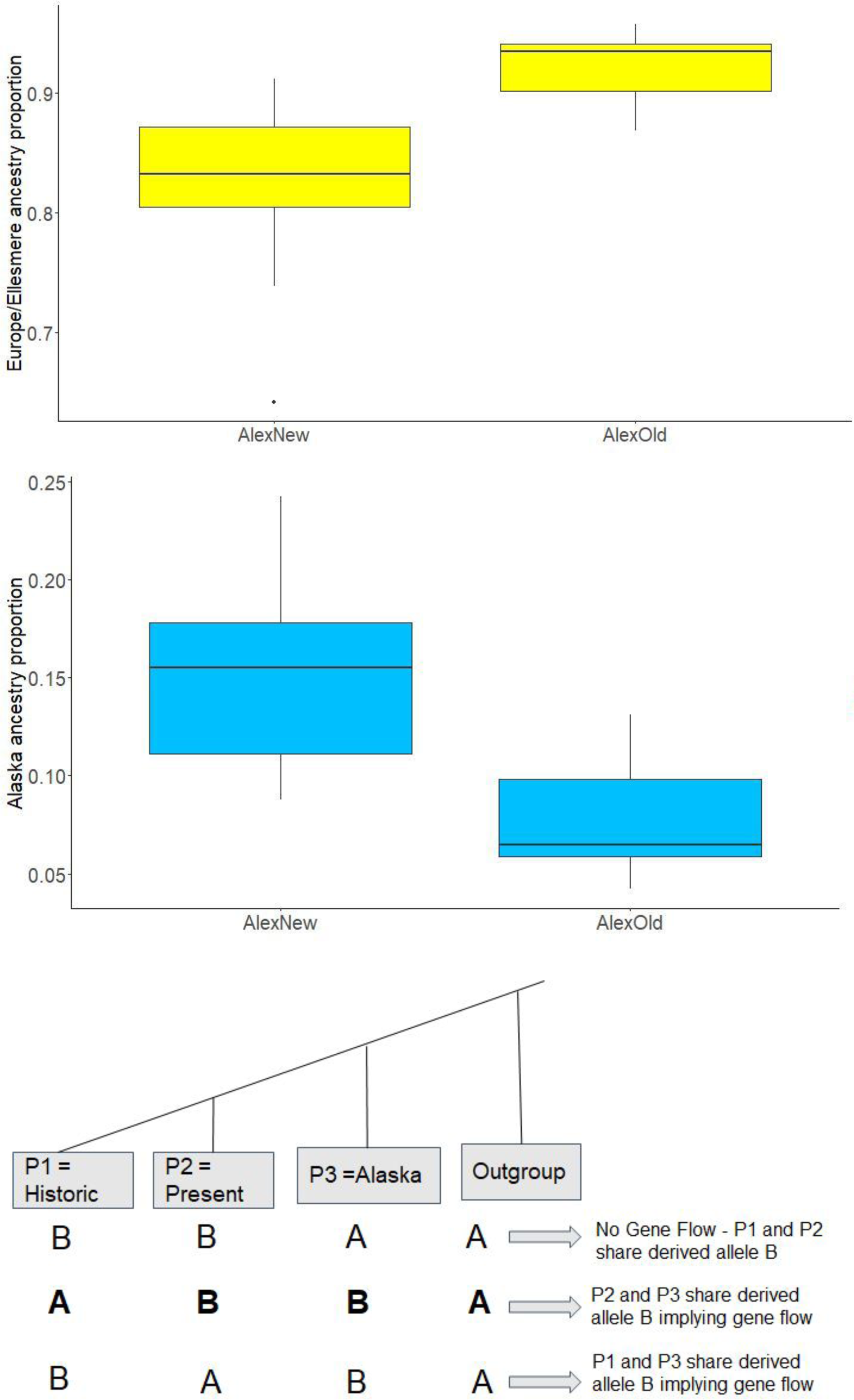
Comparison between historic and present-day samples. Samples of *C. tetragona*from Alexandra Fiord, Ellesmere Island, Canada including living individuals (AlexNew) and historic samples (AlexOld) from the ice edge of the retreating Twin Glacier. Barplots of ADMIXTURE ancestry proportions for the 23 individuals from Alexandra Fiord (based on 9285 SNPs from 330 individuals, K=5; see Fig. 1a). **(A)** Relative proportions of European ancestry observed in the present-day and historic samples. **(B)** Relative proportions of Alaskan ancestry observed in the present-day and historic samples.

## Discussion

Using GBS data from 38 populations of *C. tetragona,* including historic samples dated from the Little Ice Age, we were able to deduce the existence of multiple major glacial refugia for the Arctic in which lineages that diverged from Alaska/Beringia population 1.5 mya (Russia), 200-600 kya (Europe) and potentially 120 kya (Greenland) for *C. tetragona* survived the Pleistocene glaciations. These may also have been major refugia for other Arctic flora (Allen et al., 2015; Marr et al., 2008; Marr et al., 2013; Brochmann & Brysting, 2008). Previous studies reported that populations of *Cassiope tetragona* in Alaska have higher genetic diversity than populations found in regions glaciated during the Pleistocene, suggesting that *C. tetragona* re-colonized most of the Arctic after the last glacial maximum from Alaskan/Beringian glacial refugia (Eidesen et al., 2007). However, limited marker resolution prevented a more detailed analysis of the shallow genetic structure of the global population of *C. tetragona*. We were able to date the split between ssp. *saximontana* in southern British Columbia and ssp. *tetragona* to approximately 6 mya. In northern Canada, we observed mixing of Alaska, Europe and Greenland genetic clusters likely starting during the current interglacial (<10 kya). Moreover, using historical samples from the Little Ice Age (250-500 years old), we found that the Alaskan genotype is still spreading and expanding eastwards, as evidenced by the discovery of a higher proportion of Alaskan/foreign SNPs found in present-day Alexandra Fiord populations that are missing in the equivalent historic populations from a few hundred years ago.

### One or many refugia?

Beringia has been identified as the main refugium for Arctic plants due to its relatively high levels of intra- and inter-specific diversity (Alsos et al., 2022). Consistent with an eastwards migration out of Beringia in the last 11 kya, we found nucleotide diversity (π) was highest in Alaska and lowest in the Canadian High Arctic. We also observed a significant relationship between the date of ice retreat and the amount of diversity, similar to that found by (Pellissier et al., 2016). This suggests that recently deglaciated locations were recolonized not long after becoming ice free and/or have potentially smaller effective populations. Pollen maps have confirmed the existence of many woody trees and shrubs in Beringia at multiple times including glacial periods during the Pleistocene (Brubaker et al., 2005). A review of Arctic plant genetic clusters using AFLPs predicted that both Alaska/Beringia and Siberia were likely major refugia, but that much of North America/Europe was recolonized during the most recent interglacial from Alaska/Beringia (Eidesen et al., 2013). *Cassiope tetragona* specifically was previously assumed to have rapidly spread eastwards across North America, into Greenland and Europe recently (i.e., during the last interglacial; Eidesen et al., 2007). However, our population genomics and demographic analyses support at least three more *C. tetragona* spp. *tetragona* refugia in Europe, Greenland, and Russia, and a westward expansion of the former two. Populations in some of these refugia date back to the early Pleistocene (1.5 mya) while others formed more recently in the mid to late Pleistocene.

Many previous studies have proposed potential refugia locations based on species distribution models and past climate predictions (Pellissier et al., 2016), chloroplast DNA or AFLP data (Allen et al., 2015; Alsos et al., 2009; Eidesen et al., 2007; Eidesen et al., 2013; Marr et al., 2008; Marr et al., 2013), and pollen and fossil evidence (Brochmann & Brysting, 2008). Using chloroplast DNA data, *Bistorta vivipara* was shown to have originated in Asia and to have spread from there to Europe and Alaska before the LGM (Marr et al., 2013). Similarly, analysis of chloroplast DNA data in *Sibbaldia procumbens* found no clear evidence of a Beringia refugium, instead supporting an origin in Asia and likely refugia in Europe and southern North America (Allen et al., 2015). The strong divide observed between Europe and North America observed in *Salix herbacea* using AFLPs could also point to separate refugia (Alsos et al., 2009). Taken together, our results support previous evidence that there were regions in Europe, Alaska, and Russia that had suitable conditions for Arctic/alpine species during the glaciations.

#### Russian refugia

The Russia/Siberia cluster appears much older than the Canadian Arctic genetic clusters having diverged from other populations in the early Pleistocene (∼ 1.5 mya) possibly from Beringia/Alaska or vice versa. The divergence times that we estimated were consistent within an order of magnitude across models which used completely distinct calibration events (the outgroup divergence at 18.6 mya and the more recent range expansion into North America 7-9 kya). The relatively high F_ST_ values for the Russian populations imply little gene flow with other locations as previously noted (Eidesen et al., 2007, 2013). However, geographic sampling coverage was stronger in eastern than western Beringia, so it is possible that the Russian cluster extends eastward, and hybrid genotypes do exist in western Beringia that were not sampled in this study.

#### European refugia

European populations appear to have had their own refugia after the mid Pleistocene (divergence from Alaskan populations happened 300-600 kyrs ago). We modeled the founding of Europe from Alaska since F_ST_ values between Alaska and Europe were lower than between Russia and Europe. The decrease in European ancestry moving west into Northern Canada also supports a westward expansion from Europe after the LGM. During the current interglacial, as a result of this expansion, the European genotypes appear to have mixed with the Alaskan genotypes across Northern Canada creating some of the Canadian hybrids.

#### Greenland refugia

West Greenland/Disko Island may also have been a younger, secondary refugium that was originally populated from either the Alaskan or the European refugia. Populations in this refugium may have been founded during the late Pleistocene (∼100 kyrs), presumably during a past interglacial. This genetic cluster is found currently in high proportions in western Greenland and Baffin Island and is more differentiated from populations in Europe and Alaska than the rest of Northern Canada. “Pure” individuals for this cluster were found on Disko Island, West Greenland presenting it as a possible location where individuals may have survived the glaciations during the last 100 kyrs or been more recently introduced from an unsampled ghost population located elsewhere (Ware et al. 2012). A clear division was reported between eastern and western Greenland in Eidesen et al. (2007), which was interpreted to be a result of the large Greenland Ice Sheet. However, there is fossil evidence of *C. tetragona* in Greenland during a previous interglacial so this division may actually be a result of a secondary, young refugia in West Greenland (Bennike & Böcher, 1994). This split pattern across Greenland has been seen in several other Arctic species including *Salix herbacea* (Alsos et al. 2009, Eidesen et al., 2013). While there is also older fossil evidence of *C. tetragona* in Greenland as far back as 2 mya (Eidesen et al., 2013), our demographic results suggest that the current Greenland genetic group is younger (∼100 kyrs). This does not rule out that other genetic groups in Greenland could be unrepresented in our study or became extinct.

Two populations were sampled on Disko Island, in West Greenland (DLG and DQG) and surprisingly show entirely unique genotypes. The DQG population, only about 2.5 km away from DLG, is similar to the rest of Greenland and has relatively high F_ST_ (weighted 0.152, mean 0.110) when compared to DLG. Pollinators should transport pollen between the populations, especially from DQG to DLG, where strong easterly winds prevail. This suggests either a lack of effective gene flow between populations on the island, which could result from some strong reproductive barriers or historically separated populations that are now near each other (possibly introduced by humans from a location not sampled in this study (Ware et al. 2012)). At K=10, DLG forms its own unique ADMIXTURE cluster. We wondered if this unique population could be explained by a whole genome duplication and paralogy; however, based on flow cytometry, these populations have the same genome size.

While the main refugia were known to have had ice free regions during the glaciations (e.g. parts of Europe, Alaska, Russia and southern BC), other small refugia may have existed on nunataks in Greenland (Beatty & Provan, 2010; Westergaard et al., 2011). Although we show strong evidence for the ages of various genetic clusters and their approximate locations now, we cannot be sure of the corresponding refugium locations that would have had suitable conditions for plant survival. Based on the unique genotype on Disko Island there is a chance that a small, isolated population of *C. tetragona* survived some of the late Pleistocene there.

#### Ellesmere/Baffin divide

We observed higher F_ST_ values between Baffin Island (south) and Ellesmere Island (north) than would have been expected from their proximity. These clusters appeared to share some genetic ancestry with Greenland and Europe clusters respectively. However, they do not appear to share recent ancestry with each other, supporting the existence of a refugium in Europe and a young, secondary refugium in Greenland, as well as potential long-term barriers to gene flow between north and south Nunavut. The cause of the apparent lack of mixing between these geographically close populations is unclear, but it might be due to differences in timing of deglaciation between the Innuitian and Laurentide Ice Sheets on Ellesmere and Baffin Island, respectively (Batchelor et al., 2019). The barrier of the Greenland Ice Sheet may also have resulted in the European refugia founding Ellesmere in the north and an isolated West Greenland population founding Baffin in the south.

### Ongoing postglacial Canadian hybridization

A comparison of the Little-Ice-Age individuals (270-520 years old) with present-day populations from Ellesmere Island implies ongoing admixture between Alaskan and European genotypes in Northern Canada. Foreign SNPs appear to have been introduced to Alexandra Fiord in the last few hundred years. Based on the ADMIXTURE results at K=5 and K=10, most of these recently introduced foreign SNPs have originated in Alaska and are still mixing with the European lineage present in northern Nunavut. ABBA-BABA tests confirmed that extant populations carry more Alaskan ancestry than historical populations. Possible explanations for this ongoing mixing include (1) dominant wind currents are still present and continue to facilitate gene flow from Alaska (Hole & Macias-Fauria, 2017; Kling & Ackerly, 2021), (2) glacial retreat continues to provide space for foreign genotypes to establish (Losapio et al., 2021), (3) sexual reproduction/recombination may have been less frequent during the Little Ice Age due to shorter colder growing seasons, and (4) changing abiotic barriers continue to allow foreign genotypes to establish (Lewontin & Birch, 1966; Malcolm et al., 2002; McGraw, 1985). In addition, the increased Alaskan ancestry in Alexandra Fiord in the last few centuries suggests that the genotype from Alaska is spreading faster than the other genotypes, gradually replacing them.

To determine if missing data caused this difference between historic and present-day samples (Ewart et al., 2019), we filtered out any individuals with missing data at more than 10% of SNPs (Appendix S2 – Table S4). We observed slightly higher homozygosity in the two historic populations, which could be a result of allele dropout due to DNA degradation and more variation at restriction enzyme sites preventing DNA digestion (Andrews et al., 2016). However, this seems unlikely to completely explain the clear drop in frequency of Alaskan SNPs in the historic samples.

## Conclusions

We provide strong evidence for Pleistocene refugia in Southern BC, Beringia, Russia, Europe, and possibly a secondary, younger refugium in West Greenland. While Beringia was a refugium for North America, it appears Europe had its own unique lineage that also recolonized North America from the east during the Holocene. Multiple early to mid Pleistocene refugia for *C. tetragona* may indicate that other Arctic plant species likely also survived the glaciations in similar locations; something future studies should investigate with high resolution genomic data. The Arctic biome is logistically challenging to study and currently lacks many of the genomic resources available for many lower latitude species. However, with the climate warming at four times the global rate (Rantenan et al. 2022), Arctic species have been and will continue to be the first to deal with some major climatic shifts. Thus, it is essential to understand them from an evolutionary perspective, both to record a baseline that may be rapidly shifting as well as to help predict potential future responses in lower latitude species (Colella et al., 2020). Collaborative scientific networks, such as the International Tundra Experiment (Henry et al. 2022), allow widespread sampling and data collection for genomic studies, reducing some of the logistical constraints.

## Supporting information

S2 - Supplementary Results

## Acknowledgements

We thank Hana Christoffersen, Ed Rastetter, Gaius R. Shaver (Arctic LTER: NSF grant DEB-1637459) for sampling *Cassiope* tissue from their sites, Armando Moreno Geraldes for help and guidance regarding the GBS data analysis and Rob Elphinstone for comments regarding the paper. This research was enabled in part by WestGrid (https://www.westgrid.ca/) and Compute Canada (www.computecanada.ca). The widespread sample collection was enabled by the International Tundra Experiment network (https://www.gvsu.edu/itex/), US National Science Foundation #1504224 (to RDH), the peoples of Utqiagvik and Atqasuk, BC Parks, Canadian Museum of Nature, 2017 Students on Ice Arctic Expedition, and Canada C3 expedition. Funding for this project came from GenomeBC (sequencing), ArcticNet, Natural Sciences and Engineering Research Council of Canada (to GHRH) with logistic support from Polar Continental Shelf Program and Royal Canadian Mounted Police (to GHRH) (Ellesmere field collections), University of British Columbia, Weston Garfield Foundation, Natural Sciences and Engineering Research Council of Canada (CGSM), and Vanier (funding for CE). The Strategic Research Area: BECC - Biodiversity and Ecosystem services in a Changing Climate (to MPB and RGB) for financing the monitoring program at Latnjajaure Field Station.

## Supplementary Material

The following supplementary material is available:

**Appendix S1:** Supplementary Methods (Figure S1-S5) (Tables S1-S2) Attached

**Appendix S2:** Supplementary Results (Figures S6-S12) (Tables S3-S11) Attached

