## Supplementary material for "Multiple Pleistocene refugia for Arctic White Heather (*Cassiope tetragona*) supported by population genomics analyses of contemporary and Little-Ice-Age samples": S1- Supplementary Methods

### Appendix S1: Supplementary Methods/Protocols used

### Field Sampling Protocol

***Cassiope tetragona* sampling instructions**

**Materials**

Sampling package sent to sites that sampled:

- Enough silica for ~50 samples
- 50 empty tea bags
- 50 small Ziploc bags
- Several large Ziploc bags (some empty and others filled with the 50 small Ziploc bags containing the silica and the empty tea bags).

**Sampling Protocol**

**(Adapted from Ruud Scharn’s ITEX protocol)**

Samples from 20-50 different plants at each site were collected following the instructions below.

1. Find a *Cassiope tetragona* plant (at least a few meters from any other sampled individuals).
2. Harvest 4 fresh young *Cassiope tetragona* shoots from each plant (green leaves and the stem from the tip of the plant; Figure S1).

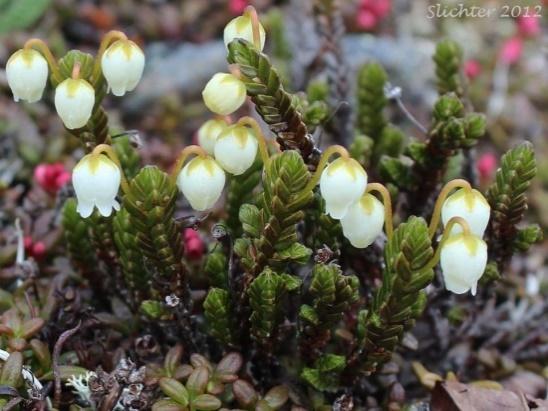

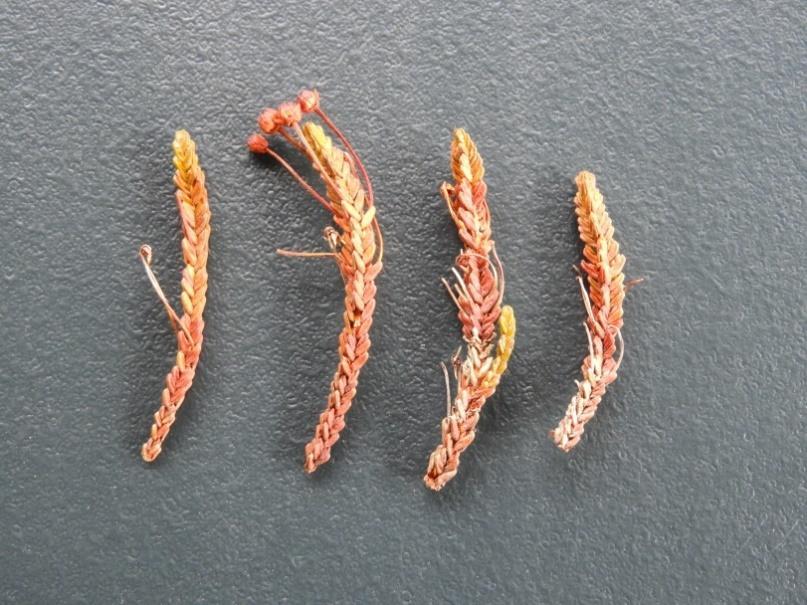

##### **Figure S1: (left)** *Cassiope tetragona* plant. **(right)** Four shoots of *C. tetragona* sampled from one plant*.*

3. Pick one of the 50 small Ziploc bags that contain silica and a tea bag from one of the large Ziploc bags.
4. Place the four shoots from one plant into the tea bag. Fold the tea bag shut.
5. Place the tea bag containing the sample back inside the small Ziploc bag filled with silica. Each plant sample should go into its own separate tea bag and small Ziploc bag (Figure S2).

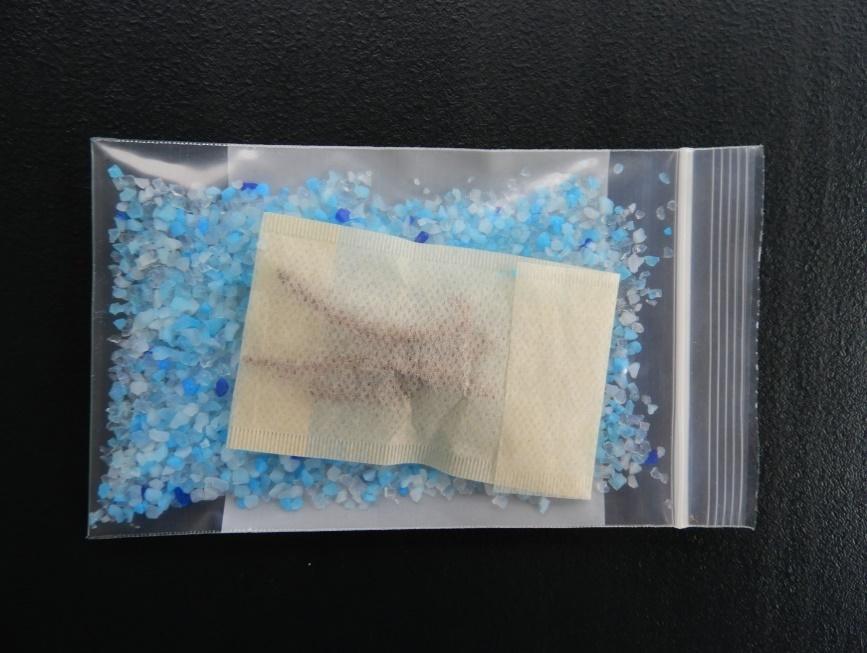

##### **Figure S2:** The four *C. tetragona* shoots placed inside the tea bag surrounded by silica in the small Ziploc bag.

6. Close the small Ziploc bag almost completely but leave a small opening for air to move through.
7. Transfer the small Ziploc bag containing the sampled plant, silica and tea bag into one of the large Ziploc bags. Extra silica (if less than 50 samples are collected) can be added to the large bags to draw out any extra moisture.
8. Continue steps 1 through 6 until all 40-50 samples have been collected. NOTE: In order to ensure all samples are collected, please keep bags containing plant samples separate from small Ziplocs containing empty tea bags.
9. Record your site’s name, latitude, longitude (Figure S3), date of collection, and name of collector on the larger Ziploc bags containing the samples. If possible, also mark each smaller Ziploc bag with an abbreviation of your site’s name.
10. Store the samples in a cool, dry, dark place until mailing. If possible, check the coloration of the leaves at regular intervals (e.g. every 2 days) for the next several days. If the leaves turn from green to brown, more silica should be added if possible.
11. Complete the sampling information form. Please try to fill in as much of the form as possible as this makes the data more valuable for future use. (Site name, number of samples collected, Date collected, Collector, GPS coordinates for the site, the approximate date of *C. tetragona* flowering at your site and an approximate average summer temperature).

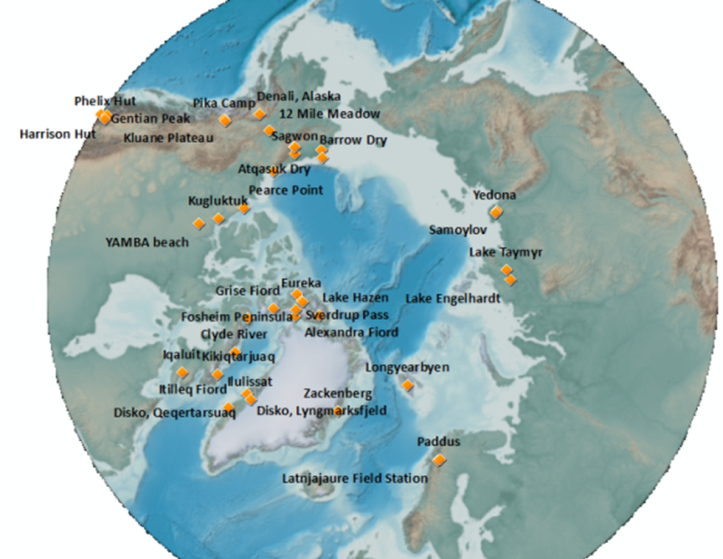

##### **Figure S3:** A map of all the sampling locations.

**Ancient sampling**

We collected the ancient samples just melting out along the base of the Twin Glacier at Alexandra Fiord and the Teardrop Glacier at Sverdrup Pass, Ellesmere Island, Nunavut, Canada. We used a rock hammer to clear away the ice in some cases at the Twin Glacier, while samples at Sverdrup Pass were collected at or within 1 m of the ice edge (Figure S4). Some samples still had green in their leaves. The plant tissue was kept frozen until it was washed free of the glacial silt and placed on silica. It was extremely wet from association with the ice, so the silica had to be repeatedly replaced.

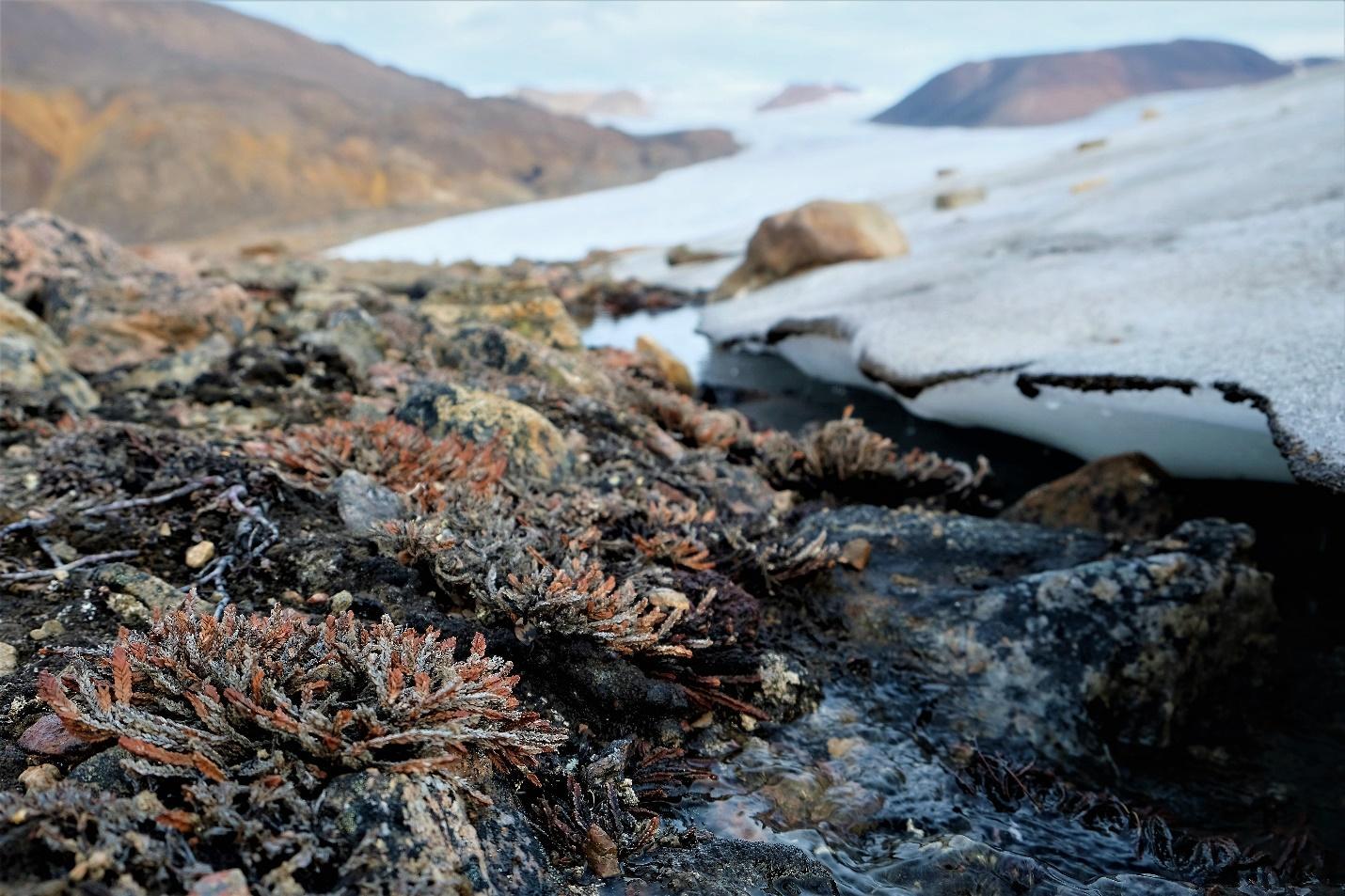

##### **Figure S4:** Over 200 year old plant communities being released from under the Twin Glacier at Alexandra Fiord.

### Modified 3% CTAB Protocol for *Cassiope tetragona*

**Modified from** **(Zeng et al., 2002)**

##### **Table S1:** 3% CTAB and buffer solutions for Arctic plants acidic leaf tissue.

| **3% CTAB solution (50 mL)** | | |  |
| --- | --- | --- | --- |
| Components | Original amounts | New for acidic tissue | Source/Catalog Number |
| Tris (1M) | 5 mL | 20 mL | Tris (1 M), pH 8.0, RNase-free  Invitrogen™/[AM9856](https://www.thermofisher.com/order/catalog/product/AM9856) |
| EDTA | 2.5 mL | 2.5 mL | Sigma-Aldrich/EDS |
| Distilled H_2_O | 30 mL | 15 mL |  |
| CTAB | 1.5 g | 1.5 g | Sigma-Aldrich/H6269 |
| 3% PVP | 0.5 g | 1.5 g | Sigma-Aldrich/PVP40 |
| NaCl | 4.38 g | 4.38 g |  |
| top up to 50 mL at the end | | |  |
| **CTAB Free Buffer (50 mL )** | | |  |
| Components | Original amounts | New for acidic tissue |  |
| Tris (1M) | 10 mL | 20 mL | Tris (1 M), pH 8.0, RNase-free  Invitrogen™/[AM9856](https://www.thermofisher.com/order/catalog/product/AM9856) |
| EDTA | 5 mL | 5 mL | Sigma-Aldrich/EDS |
| H_2_O | 25 mL | 15 mL |  |
| NaCl | 0.73 g | 0.73 g |  |
| 3% PVP | NA | 1.5 g | Sigma-Aldrich/PVP40 |
| top up to 50 mL at the end | | |  |

1.	Cool #samples x 1 mL CTAB-free buffer + #samples x 6 µL β-mercaptoethanol

2.	Warm #samples x 500 μL 3% CTAB + #samples x 5 μL of β-mercaptoethanol

3.	Grind frozen tissue, 2 min in grinder, try liquid nitrogen

4.	Add 1 mL of CTAB-free buffer (cold) + 6 µL β-mercaptoethanol

5.	Mix by inversion

6.	Keep 10 minutes on ice

7.	Spin at 10,000 g for 10 minutes

8.	Discard the supernatant

9.	Add 500 μL of pre-warmed 3% CTAB + 5 μL of β-mercaptoethanol

10.	Add 3 μL of RNAseA (20 mg/ml)

11.	Incubate at 65°C for 60-120 minutes in a water bath

12.	Let the tubes cool down a little bit

13.	Add one volume (500 μL) of phenol: chloroform: isoamyl alcohol (25:24:1) to your sample, and vortex or shake by hand thoroughly for approximately 20 seconds.

14.	Centrifuge at room temperature for 5 minutes at 16,000 g.

15.	Carefully remove the upper aqueous phase and transfer the layer to a fresh tube. Be sure not to carry over any phenol during pipetting.

16.	Add 0.1 volumes of NaCl 5M, mix

17.	Add 0.7 volumes of cold isopropanol, mix

18.	Leave 30 minutes at -20°C

19.	Spin at max speed for 15 minutes

20.	Wash pellet with 1 mL of cold 75% ethanol

21.	Spin at max speed for 5 minutes

22.	Repeat wash

23.	Dry pellet (incubate for 15 minutes in 37°C)

24.	Resuspend pellet in 50 μL of 10 mM Tris pH 8.0

### Genotyping by sequencing library preparation protocol

Based off of the Rieseberg lab blog: <https://www.zoology.ubc.ca/~rieseberg/RiesebergResources/wp-content/uploads/2018/02/Simplified-GBS-protocol-2017.pdf>
Samples were randomized in the plates and were each assigned a unique combination of in-line barcodes on both the PE1 adapter (New England Biolabs Restriction enzyme PstI Hi-Fidelity; 192 unique barcodes) and the PE2 adapter (New England Biolabs Restriction enzyme MspI, 12 unique barcodes) (Andrews et al., 2016). To improve uniformity of representation across all individuals (especially for degraded ancient DNA samples), samples were quantified and pooled after the PCR enrichment step. Each plate of samples was pooled together, and the ancient samples were pooled separately so they could be added to the final libraries at a higher concentration. Samples were only included in the pool if they had successful PCR amplification concentrations. After pooling, fragments in the 300-500 bp range were gel size-selected from each pooled plate and the ancient samples.

The GBS library preparation for the ancient samples was identical to the present-day samples except that at the pooling stage we included twice as much DNA than what was used for present-day samples; this was done because we expected a lower proportion of reads from ancient samples to be informative, due to the lower initial DNA quality. Over time, DNA degrades because of oxidation and hydrolysis (Gugerli et al., 2005). Using an agarose gel, we analyzed the length of DNA fragments from the ancient *C. tetragona* samples. DNA fragments were found that ranged in length from 100-5000 bp for all the samples selected for GBS. After sequencing, we used FastQC (Andrews, 2010) to compare the quality of ancient and present-day samples.

To minimize the presence of high-copy fragments deriving from the chloroplast and mitochondria genomes, we performed an enzymatic depletion of repetitive sequences (Matvienko et al., 2013; Moyers et al., 2017; Todesco et al., n.d.). The fragment size distribution in the completed libraries was examined using a Bioanalyzer high-sensitivity DNA chip (Agilent).

The detailed depletion protocol can be found on the Rieseberg Lab blog: <https://www.zoology.ubc.ca/~rieseberg/RiesebergResources/wp-content/uploads/2015/06/Depletion-of-repetitive-sequences-GBS-libraries.pdf>

### Flow cytometry protocol and results

**Preparation**

- Make filters, 2 layers of miracloth between plastic pieces
- Cut tips for 1mL pipette. Make one per sample.
- Label a 2mL for each sample/standard combination

##### **Table S2:** Nuclei extraction and wash buffer was slightly modified from De Laat's buffer (de Laat and Blaas, 1984) and based off of the Rieseberg Lab blog: <https://www.zoology.ubc.ca/~rieseberg/RiesebergResources/dna-ploidy-and-genome-size-estimation-using-flow-cytometry/>

| **For 250mL buffer** | **Amount** |
| --- | --- |
| 0.894g | 15 mM HEPES |
| 0.5mL of 0.5M | 1mM EDTA |
| 0.5mL | 0.2 % (v/v) Triton X-100 |
| 1.49g | 80 mM KCl |
| 0.292g | 20 mM NaCl |
| 25.7g | 300 mM sucrose |
| 12.5mL | 0.5 mM spermine |
| 0.262mL – add after filtering | 15 mM β-mercaptoethanol |
| 2.5g | 0.25 mM PVP-40 |

Add all the above to about 150mL.
Adjust volume to 250mL and the pH of this buffer to 7.
Filter out contaminants
Add 2-mercaptoethanol at the end.

**Steps**

1)	Measure out 30-40mg of leaf tissue into a petri dish for each of standard+standard, standard+sample and sample+sample. Keep tissue in the fridge, moist while waiting to cut.

2)	Chop up the tissue rapidly in 1mL of cold buffer on a cold aluminum block. Cut rapidly for approximately 1 minute.

3)	Using the cut tips, move buffer and tissue to the double miracloth filter placed in a labeled 2mL tube. Wait for all the liquid to flow through.

4)	Spin at room temperature @ 1200 RPM (SLOW!) for 5min. After this you should see a white (or often green) pellet. Carefully pipette out the supernatant. The pellet may not be tightly bound to the wall of the tube.

5)	Resuspend pellet in 1mL of buffer and repeat Step 4 above.

6)	Fix nuclei in ethanol:acetic acid (v:v) 3:1. Mix well by inversion and store at 4 degrees Celsius overnight.

7)	Repeat Step 4. Can do an extra spin @1500 RPM for 1min if pellet is not sticking to the wall.

8)	Add 200mL of buffer.

9)	Add 4uL of 20mg/mL RNAase A. Wait approx.. 30 min at room temp.

10) Add the propidium idodide right before leaving to go to the flow cytometer. Add 20uL to the 200mL of buffer. The more dye added the wider the peaks may be.

**Results**

From the fluorescence peaks we calculate the genome size:
Sample_2C _pg = (Standard_2C_pg * mean_sample_peak)/ mean_standard_peak

Conversion C (pg) to base pairs:
Diploid_bp = 0.978 *10^9^ * Sample_2C_pg

**Standard plants’ seeds in the lab:**

- *Raphanus sativus* (2C = 1.11 pg DNA)
- *Solanum lycopersicum* (2C = 1.96 pg DNA)
- *Glycine max* (2C = 2.50 pg DNA)
- *Zea mays* (2C = 5.43 pg DNA)
- *Pisum sativum* (2C = 9.09 pg DNA)
- *Secale cereale* (2C = 16.19 pg DNA)
- *Vicia faba* (2C = 26.90 pg DNA)

**Seeds received November 2021 from:**
Prof. Ing. Jaroslav Doležel, DrSc.
Centrum strukturní a funkční genomiky rostlin
Ústav experimentální botaniky AV ČR
Šlechtitelů 31
783 71 Olomouc

Web: <http://olomouc.ueb.cas.cz>

### Demographic models

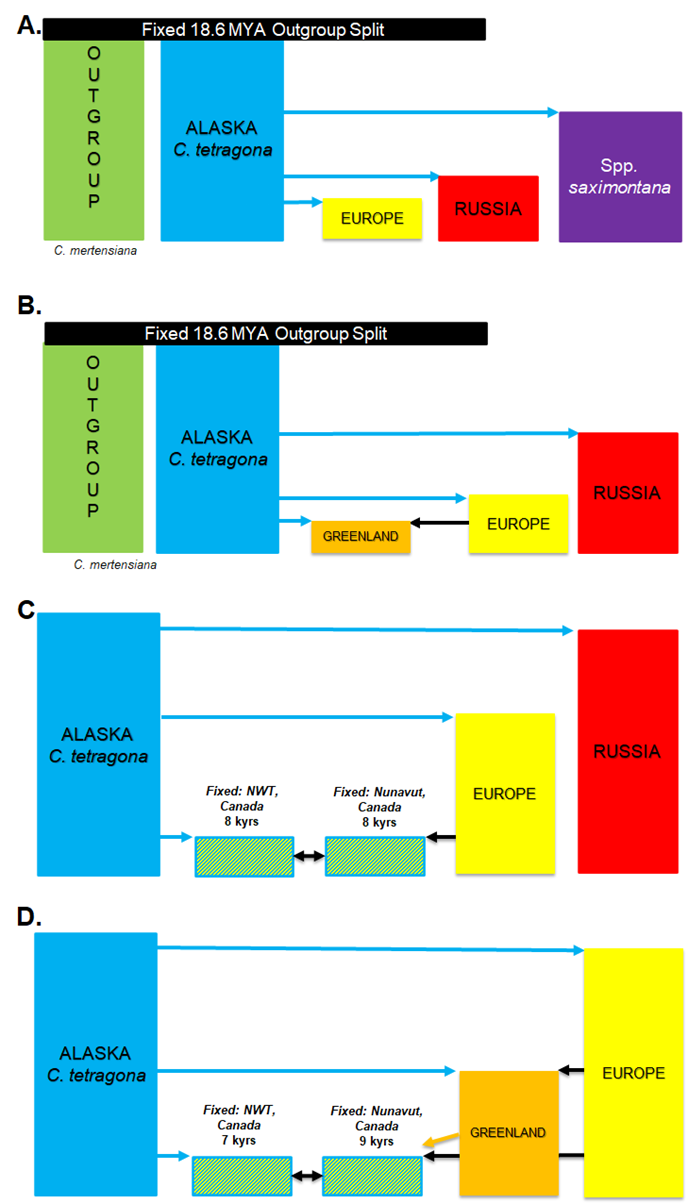

##### **Figure S5:** Schematic of fastsimcoal2 demographic models used to estimate refugia ages. ModelA: *Cassiope* *mertensiana*, *C. tetragona* ssp. *saximontana*, Russia, Alaska and Europe, ModelB: *C. mertensiana*, Russia, Alaska, Europe and Greenland, ModelC: Russia, Alaska, Europe, NWT and Nunavut, and ModelD: Alaska, Europe, Greenland, NWT and Nunavut

### Literature Cited

###### de Laat, A. M., & Blaas, J. (1984). Flow-cytometric characterization and sorting of plant chromosomes. *TAG. Theoretical and Applied Genetics. Theoretische Und Angewandte Genetik*, *67*(5), 463–467. https://doi.org/10.1007/BF00263414
