## Supplementary material for "Multiple Pleistocene refugia for Arctic White Heather (*Cassiope tetragona*) supported by population genomics analyses of contemporary and Little-Ice-Age samples": S2 - Supplementary Results

### Appendix S2: Supplementary results

### Sample information

##### **Table S3:** Sample information for 349 *Cassiope* plants included in analyses. Plants correspond to *Cassiope tetragona* ssp. *tetragona* unless otherwise specified. Populations are designated with short population codes (column1). Colours represent the ADMIXTURE cluster at K=5 (and *C. mertensiana*, used as the outgroup in analyses) that made up the largest proportion of that population’s genotypes (Green = *C. mertensiana*; purple = *C. tetragona* ssp. *saximontana*; blue = Alaska; orange = Greenland; yellow = Europe; and red = Russia).

| **Population code** | **Site** | **Region** | **Collector** | **Date of sample collection (2017)** | **# samples sequenced** | **Latitude** | **Longitude** | **Elevation (m)** | **Deglaciation date (thousand years BP)** |
| --- | --- | --- | --- | --- | --- | --- | --- | --- | --- |
| MER | *C. mertensiana* Harrison Hut (HAR), Phelix Hut (PHE) | British Columbia | Cassandra Elphinstone | June 18 , Oct 29 | 9 | 50.5459 | -123.2652 | 1689 | NA |
| GEN | Gentian Peak (*C. tetragona* ssp. *saximontana*) | British Columbia | Cassandra Elphinstone | Aug 19 | 10 | 49.9511 | -122.99 | 2197 | NA |
| ATQ | Atqasuk Dry | Alaska | Robert Hollister, Jacob Harris, Hana L. Christoffersen | June 24 | 10 | 70.4537 | -157.4074 | 22 | NA |
| BARD | Barrow (Utqiaġvik) Dry | Alaska | Robert Hollister, Jacob Harris, Katlyn R. Betway, Hana L. Christoffersen | July 8 | 10 | 71.315 | -156.5989 | 5 | NA |
| DEN | Denali | Alaska | Michael Moody | July 31 | 10 | 63.0769 | -146.223 | 1022 | 12.5 |
| IMN | Imnavait Creek | Alaska | Ed Rustetter, Erin Gleeson, Gus Shaver | July 26 | 10 | 68.6174 | -149.3092 | 925 | NA |
| MAT | Toolik Lake Moist Acidic | Alaska | Ed Rustetter, Erin Gleeson, Gus Shaver | July 26 | 9 | 68.6344 | -149.6433 | 753 | 12 |
| MIL | 12 Mile Meadow | Alaska | Michael Moody | July 29 | 10 | 65.3975 | -145.973 | 962 | NA |
| MNT | Toolik Lake Moist Non-Acidic | Alaska | Ed Rustetter, Erin Gleeson, Gus Shaver | July 26 | 10 | 68.6236 | -149.6103 | 760 | NA |
| SAG | Sagwon | Alaska | Michael Moody | July 25 | 10 | 69.4244 | -148.6976 | 297 | NA |
| KL | Kluane Lake (West) | Yukon (Kluane) | Haydn Thomas | Exact date unknown (2017) | 10 | 60.9811 | -138.4113 | 1397 | 12 |
| PC | Kluane Lake (East) | Yukon (Kluane) | Haydn Thomas | Exact date unknown (2017) | 9 | 61.2114 | -138.2845 | 1710 | 13 |
| QHI | Qikiqtaruk | Yukon | Isla Myers-Smith, Gergana Daskalova | July 30 | 8 | 69.5789 | -138.9114 | 48 | 15 |
| PEA | Pearce Point | NWT | Paul Sokoloff | Sept 6 | 9 | 69.8223 | -122.6625 | 0-10 | 9.5 |
| YAM | YAMBA, Daring Lake | NWT | Wan Case, Joachin Obst, Karin Clark | Aug 10 | 9 | 65.8667 | -111.5333 | 450 | 9.5 |
| AXE | Axel Heiberg Island | Nunavut | Paul Sokoloff, Troy McMullin | July 12 | 9 | 79.4151 | -90.7481 | 200 | 2 |
| AlexNew | Alexandra Fiord | Ellesmere, Nunavut | Cassandra Elphinstone, Sofie Agger, Katie MacIntosh, Jamie Blackley, Greg Henry | July 7-31 | 14 | 78.8648 | -75.8007 | 16 | 7.2 |
| AlexOld | Alexandra Fiord, Historic, Twin Glacier | Ellesmere, Nunavut | Cassandra Elphinstone, Sofie Agger, Katie MacIntosh, Jamie Blackley, Greg Henry | July 31 | 9 | 78.8452 | -75.7996 | 100 | 7.2 |
| SVN | Sverdrup Pass | Ellesmere, Nunavut | Cassandra Elphinstone, Sofie Agger, Katie MacIntosh, Jamie Blackley, Greg Henry | July 17 | 1 | 79.1167 | -80.6667 |  | 7.8 |
| SVO | Sverdrup,  Historic, Teardrop Glacier | Ellesmere, Nunavut | Cassandra Elphinstone, Sofie Agger, Katie MacIntosh, Jamie Blackley, Greg Henry | July 17 | 4 | 79.1167 | -80.6667 |  | 7.8 |
| FOS | Combined Eureka, Fosheim Penninsula | Ellesmere, Nunavut | Paul Sokoloff, Troy McMullin | July 7-11 | 8 | 80.0593 | -85.3122 | 75-230 | 8.5 |
| GF | Grise Fiord | Ellesmere, Nunavut | Cassandra Elphinstone | August 10 | 3 | 76.4261 | -82.9094 |  | 9.5 |
| HAZ | Lake Hazen | Ellesmere, Nunavut | Paul Sokoloff, Troy McMullin | July 16 | 10 | 81.8274 | -71.3366 | 250 | 5.5 |
| CR | Clyde River | Baffin, Nunavut | Cassandra Elphinstone | July 2 | 10 | 70.4844 | -68.5131 | 27 | 10.5 |
| BY | Bylot Island | Baffin, Nunavut | Esther Lévesque and Vincent Maire | July 11 | 10 | 73.1569 | -79.9687 | 20 | 10.5 |
| Iq | Iqaluit | Baffin, Nunavut | Greg Henry | Sept 6 | 9 | 63.7467 | -68.517 |  | 8.5 |
| Kik | Qikiqtarjuaq | Baffin, Nunavut | Greg Henry | Sept 1 | 10 | 67.5556 | -64.0257 |  | 13 |
| KUQ | Kugluktuk | Baffin, Nunavut | Paul Sokoloff | Sept 3 | 6 | 67.7984 | -115.2317 | 35 | 10.5 |
| 25 | Itilleq Fiord | Greenland | Roger Bull | Aug 20 | 10 | 66.5326 | -53.4156 | 20 | 11.5 |
| IG | Ilulissat | Greenland | Per Molgaard | June 26 | 9 | 69.207 | -51.105 | 55 | 9.5 |
| DQG | Qeqertarsuaq | Disko, Greenland | Per Mølgaard | July 1 | 10 | 69.2541 | -53.5176 | 20 | 11.5 |
| DLG | Lyngmarksfjeld | Disko, Greenland | Per Mølgaard | June 29 | 10 | 69.2703 | -53.5635 | 390 | 11.5 |
| ZAC | Zackenberg | Greenland | Niels Martin Schmidt | July 7 | 10 | 74.47 | -20.5617 | 75 | 10.5 |
| PET | Petunia Bay | Svalbard | Esther Frei | July 27 | 5 | 78.711112 | 16.45888 |  | 15 |
| LAJ | Latnjajaure | Sweden | Mats P. Björkman, Robert G. Björk | July 23-24 | 10 | 68.35 | 18.5 |  | 10 |
| LON | Longyearbyen | Svalbard | Esther Frei | July 27 | 10 | 78.207572 | 15.606609 |  | 13 |
| SW | Paddus | Sweden | Annika Hofgaard | Sept 18 | 10 | 68.3247 | 18.9583 | 550 | 9 |
| YED | Kurunaghk, Yedoma, Lena River Delta | Russia | Julia Boike | Sept 9 | 10 | 72.36811 | 126.25771 Kurunaghk is part of of the Yedoma ice rich pleistocene landforms. |  | NA |
| SAM | Samoylov, Lena River Delta | Russia | Julia Boike | Sept 12 | 9 | 72.3772 | 126.4792 |  | NA |

### SNP Filtering

##### **Table S4:** Amount of missing data for each individual. The population code following “Pop” (e.g. 025 or AlexNew) can be found referenced in Table 1 in the main paper. <https://github.com/celphin/Population_genomics_Cassiope/blob/main/Figures_data/Missing_data_table.csv>

##### **Table S5:** **Percentage of eukaryotic and bacterial reads in the historic samples from under the ice. The reads were BLASTed to the NCBI database and identified to Kingdom.** The percent of reads that mapped to the de novo reference was slightly lower than the eukaryotic read percentage for each individual. SVO = Sverdrup, Teardrop Glacier; AlexOld = Alexandra Fiord, Twin Glacier.

| **Sample** | **% eukaryotic reads** | **% bacterial reads** |
| --- | --- | --- |
| PopSVO_20.R1 | 2.9 | 97.1 |
| PopSVO_20.R2 | 3.3 | 96.7 |
| PopSVO_1.R2 | 4.1 | 95.9 |
| PopSVO_13.R1 | 4.4 | 95.6 |
| PopSVO_18.R1 | 4.6 | 95.4 |
| PopSVO_18.R2 | 4.6 | 95.4 |
| PopSVO_1.R1 | 5.2 | 94.8 |
| PopSVO_13.R2 | 5.3 | 94.7 |
| PopSVO_15.R1 | 5.4 | 94.6 |
| PopSVO_15.R2 | 5.6 | 94.4 |
| PopSVO_28.R2 | 5.8 | 94.2 |
| PopSVO_28.R1 | 6.1 | 93.9 |
| PopAlexOld_33.R2 | 6.3 | 93.7 |
| PopAlexOld_66.R1 | 6.6 | 93.4 |
| PopSVO_16.R2 | 7.6 | 92.4 |
| PopAlexOld_66.R2 | 8.0 | 92.0 |
| PopSVO_16.R1 | 8.0 | 92.0 |
| PopSVO_5.R1 | 8.0 | 92.0 |
| PopAlexOld_33.R1 | 8.1 | 91.9 |
| PopSVO_5.R2 | 10.4 | 89.6 |
| PopSVO_36.R2 | 13.8 | 86.2 |
| PopAlexOld_21.R2 | 14.2 | 85.8 |
| PopAlexOld_21.R1 | 15.1 | 84.9 |
| PopSVO_38.R2 | 16.8 | 83.2 |
| PopSVO_38.R1 | 17.5 | 82.5 |
| PopSVO_24.R2 | 17.9 | 82.1 |
| PopAlexOld_56.R2 | 18.4 | 81.6 |
| PopSVO_37.R1 | 19.1 | 80.9 |
| PopSVO_37.R2 | 20.4 | 79.6 |
| PopSVO_24.R1 | 22.4 | 77.6 |
| PopSVO_36.R1 | 22.6 | 77.4 |
| PopAlexOld_56.R1 | 23.6 | 76.4 |
| PopAlexOld_50.R2 | 24.7 | 75.3 |
| PopAlexOld_24.R2 | 30.7 | 69.3 |
| PopAlexOld_24.R1 | 33.3 | 66.7 |
| PopAlexOld_30.R2 | 40.4 | 59.6 |
| PopAlexOld_30.R1 | 42.8 | 57.2 |
| PopAlexOld_89.R2 | 62.5 | 37.5 |
| PopAlexOld_89.R1 | 63.3 | 36.7 |

### Principal Component Analysis

##### **Table S6: Principal component analysis** percent variance based on data with MAC filtering of 2 and MAF filtering of 5%.

**MAF=1**

PC1 = 27.00

PC2 = 5.62

PC3 = 2.63

PC4 = 1.63

PC5 = 1.50

PC6 = 1.26

PC7 = 1.02

PC8 = 0.93

PC9 =	0.93

PC10 = 0.85

### ADMIXTURE

##### **Table S7: ADMIXTURE 1.3.0** cross validation errors for K values between 1 and 14 based on 330 individuals and 9 285 SNPs.

| **K value** | **Cross Validation Error (CV) Take 1** | **Cross Validation Error (CV) Take 2** | **Cross Validation Error (CV) Take 3** |
| --- | --- | --- | --- |
| 1 | 0.36984 | 0.36982 | 0.36980 |
| 2 | 0.30263 | 0.30255 | 0.30261 |
| 3 | 0.28295 | 0.28303 | 0.28300 |
| 4 | 0.26980 | 0.26979 | 0.26959 |
| **5** | **0.26186** | **0.26204** | **0.26197** |
| 6 | 0.25857 | 0.25863 | 0.25829 |
| 7 | 0.25699 | 0.25659 | 0.25681 |
| 8 | 0.25598 | 0.25458 | 0.25547 |
| 9 | 0.25239 | 0.25220 | 0.25599 |
| 10 | 0.25155 | 0.25522 | 0.25562 |
| 11 | 0.25111 | 0.25087 | 0.25249 |
| 12 | 0.25299 | 0.25212 | 0.25105 |
| 13 | 0.25106 | 0.25070 | 0.25129 |
| 14 | 0.25303 | 0.25135 | 0.25259 |

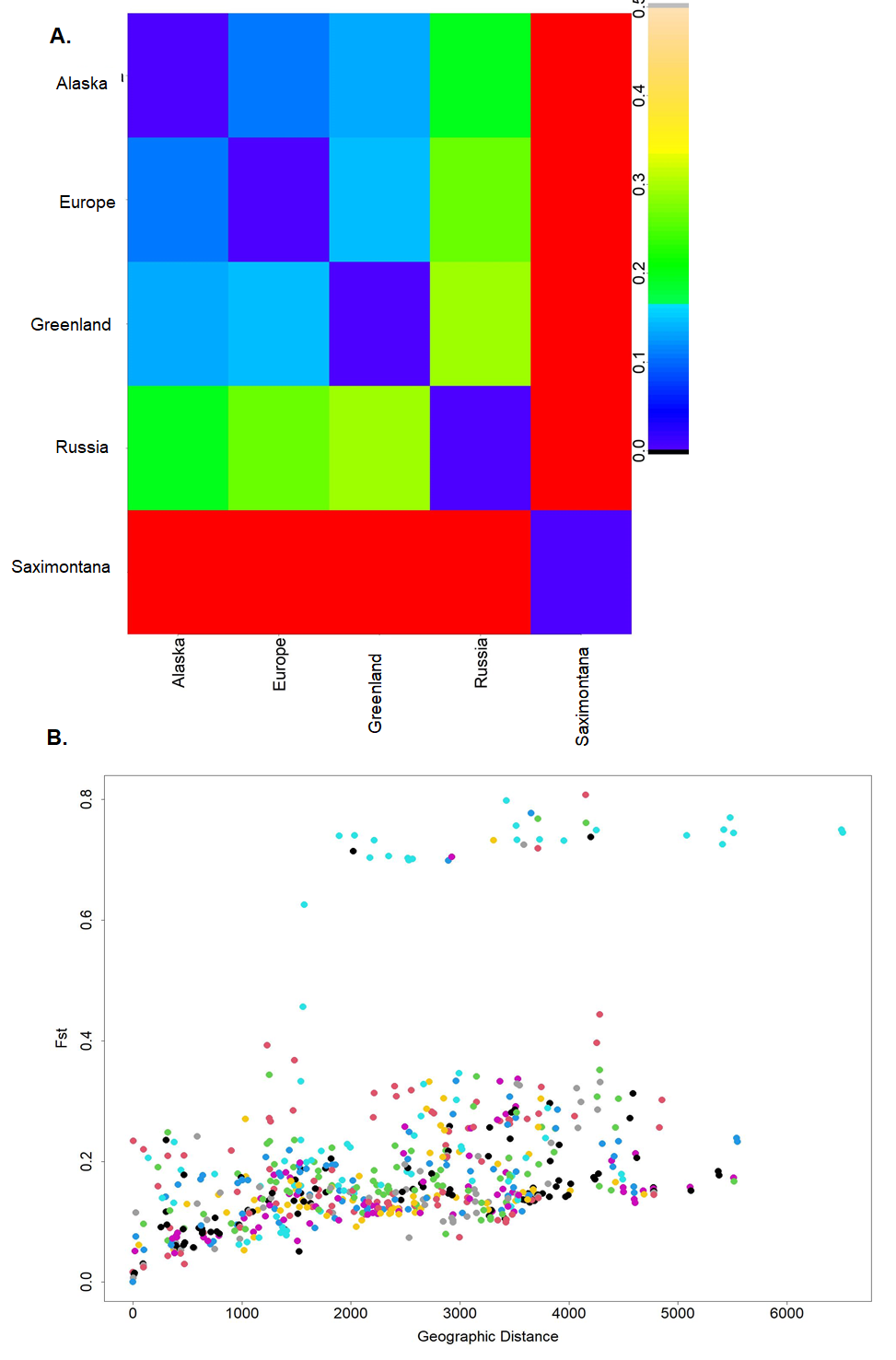

##### **Figure S6: F_ST_ results** based on 9 285 SNPs from 330 individuals in ADMIXTURE. **(A)** ADMIXTURE clusters F_ST_ plot using 5 groups (CV error=0.158). F_ST_ values >0.3 are shown in red (outgroup = MER and British Columbia *saximontana* population = GEN), missing data in grey and =<0 in black. **(B)** Mantel test plot: genetic distance F_ST_ versus ln(geographic distance(km)) (Mantel test, p-value=2 e^-4^).

### Population statistics (Tajima’s D, heterozygosity), abiotic factors and leaf weights

##### **Table S8:** For each population average population statistics (nucleotide diversity, Tajima’s D, inbreeding coefficient), ADMIXTURE proportions, and principal component amounts are shown.

<https://docs.google.com/spreadsheets/d/1SqkSMOfKzwpvAxs0X5yc78JY8Tw7Ox7ObgrvzwmNX7M/edit?usp=sharing>

The inbreeding coefficient, F_IS_ ($\frac{H_{e}-H_{o}}{H_{e}}$) was calculated for each individual based on the allele frequencies within its population. To quantify heterozygosity, F_IS_ was determined for each individual from the total number of segregating sites and the observed/expected homozygosity (population averages shown in Table S8). Arctic populations frequently had more outbred individuals than expected by HWE (F_IS_<0, typically around -0.3; Figure 3B). The BC *C. tetragona* ssp. *saximontana* subspecies (GEN; F_IS_ =-0.47), as well as the population in Disko, Greenland (DLG: F_IS_ =-0.62), were highly heterozygous, but exhibited very low genetic diversity (GEN: π =8.1, DLG: π =12, compared to π =~20 for most other populations ). The historic population (AlexOld, Table 1) appeared to be closer to HWE (F_IS_=0).

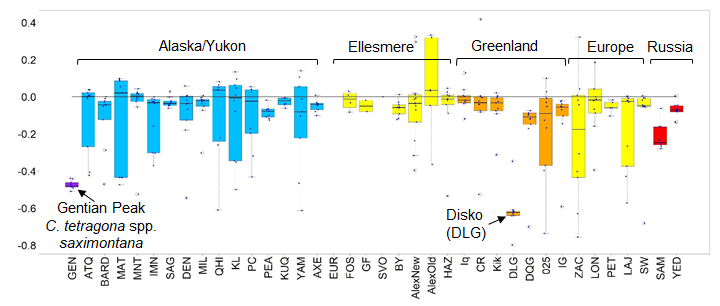

##### **Figure S7: Inbreeding coefficient**. Boxplots of F_IS_ (inbreeding coefficient) for populations sorted by geographic location (total variant sites=26 350) (ordered alphabetically by region). Most Arctic populations show slightly higher than expected heterozygosity (F_IS_<0). Zero line is shown. Gentian Peak, BC (GEN) and Disko Island, Greenland (DLG) show extremely high outbreeding possibly due to poor mapping to the reference genome for these groups (resulting in higher than expected heterozygosity).

Tajima’s D, a measure of the frequency of rare alleles in populations, was calculated across windows of 150 SNPs/variants in vcftools for each population and then all the window values were averaged for each population. Positive values indicate fewer rare alleles and negative ones indicate many rare alleles. The average Tajima’s D was consistently slightly positive for most populations implying few rare alleles were found in most locations (Table E-1).

Even though the population on Gentian Peak in southern BC, Canada appeared to be quite small (only roughly 20-30 individuals), it was more heterozygous when compared to HWE than any other population measured. This could imply that there is gene flow from other locations reaching this population even though it appears geographically isolated. Although the slightly positive Tajima’s D observed in most of the populations is likely a result of ascertainment bias favouring common alleles and missing some rare alleles, the relatively higher Tajima’s D (very few low frequency alleles) for Gentian Peak and small population size may indicate a bottleneck event in the past. The neighbor-net tree indicated individuals within this population were more closely related to each other than in many Arctic populations perhaps indicating all individuals may have recently expanded from a single colonization event although this does not explain their high heterozygosity. It is possible that higher than expected heterozygosity was observed due to poor mapping of this more distant subspecies’ reads onto the *de novo* reference. Establishment outside the ‘good’ growing conditions on Gentian Peak, appears to be quite difficult since there are no other populations in the vicinity.

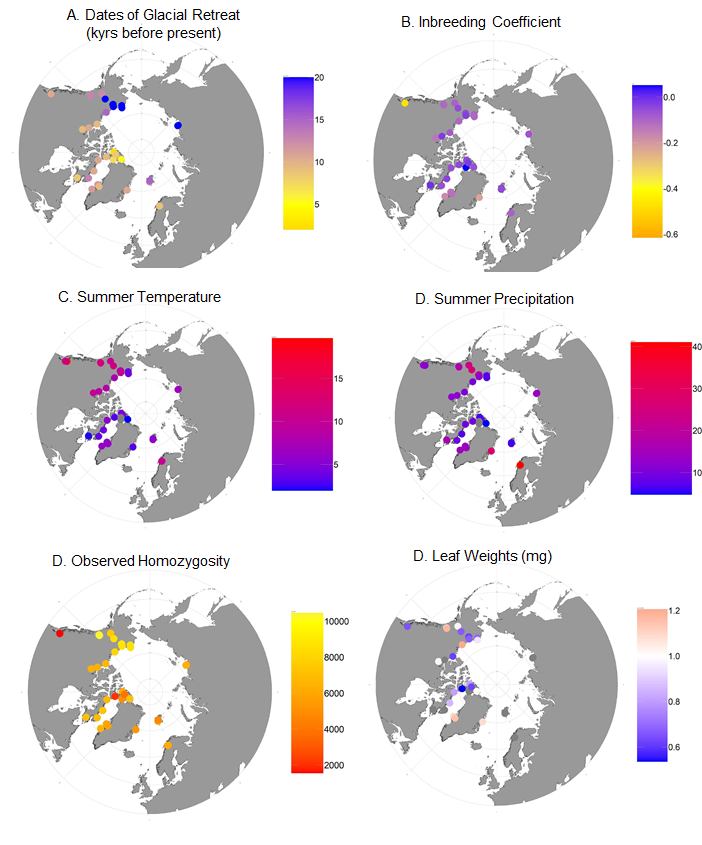

##### **Figure S8: Maps showing geographic patterns**. (A) Date of deglaciation after the last major glacial period in thousands of years ago at each of the sampling sites. Dates from (Dalton et al., 2020; Hughes et al., 2016). (B) Tajima’s D plotted for 150bp windows across the genome. (C) Map showing the estimated average precipitation from the nearest climate station for each of the sampled sites. (D) Map of average summer (June-August) temperatures from 1997-2017 from the nearest climate station to the sampling location. (E) Observed homozygosity. (F) Average leaf weight measures (mg) at each sampling site. Green leaves (20) were removed from each plant, dried on silica, and then weighed. The weight was divided by 20 for each plant. Between 2 and 5 plants were measured at each site shown. Grey sites had no data. Leaf weights were correlated with climate data but no significant result was found.

### TreeMix Results

###### TreeMix 1.13 (Pickrell & Pritchard, 2012) was used to find the maximum likelihood tree showing the relationship between the 6 main groups identified by ADMIXTURE (K=5 plus the outgroup). For this, each individual was assigned to their dominant ADMIXTURE cluster (membership >0.5) and then a tree was built showing the relationship between the groups. For each tree, edges (gene flow) from 0 to 5 were included. SNPs missing data in any individual were filtered prior to the analysis.

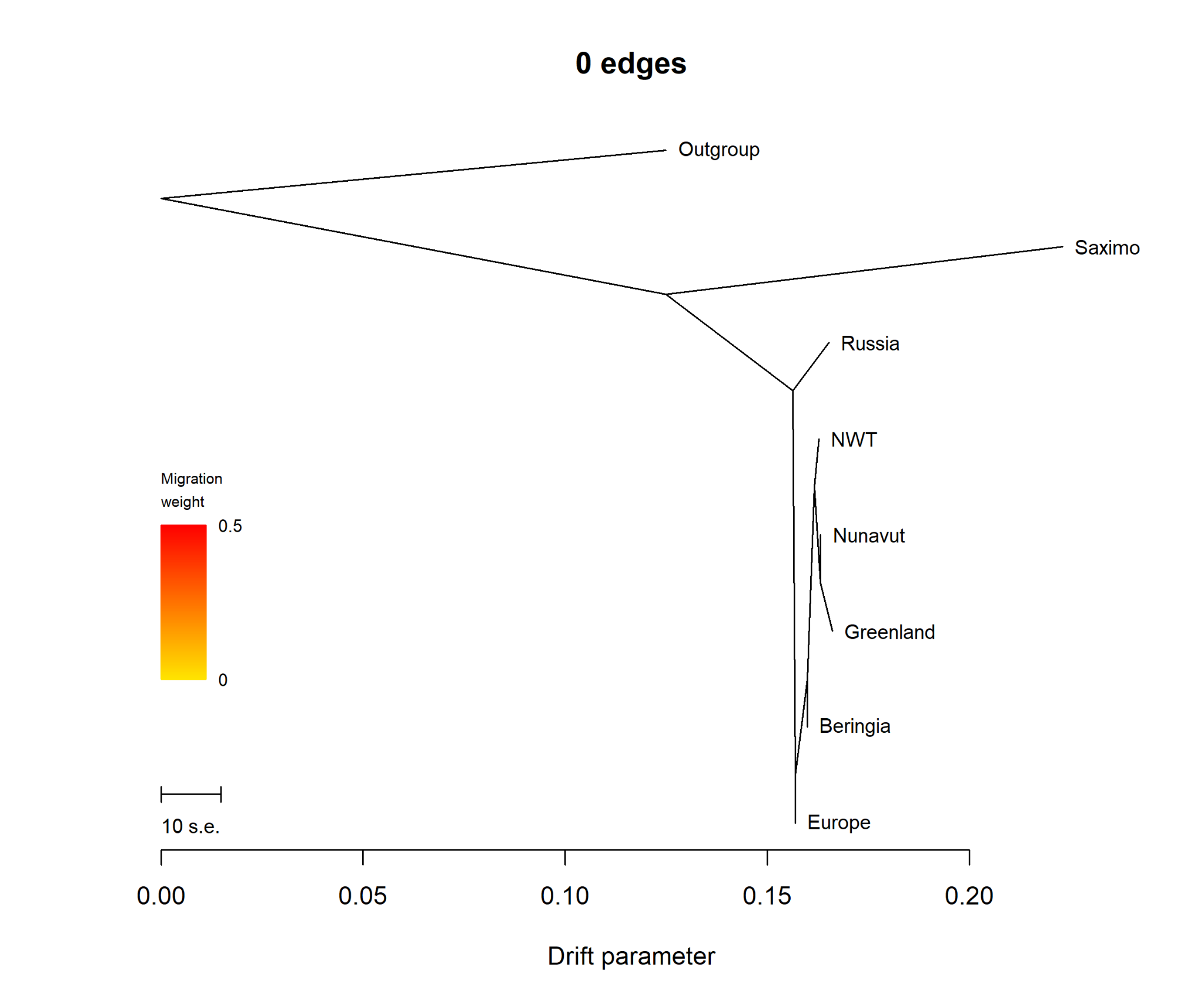

##### **Figure S9:** **Maximum likelihood tree made using 349 individuals and 2310 SNPs in TreeMix 1.3.0**. Populations were grouped by their dominant cluster in ADMIXTURE K=6.

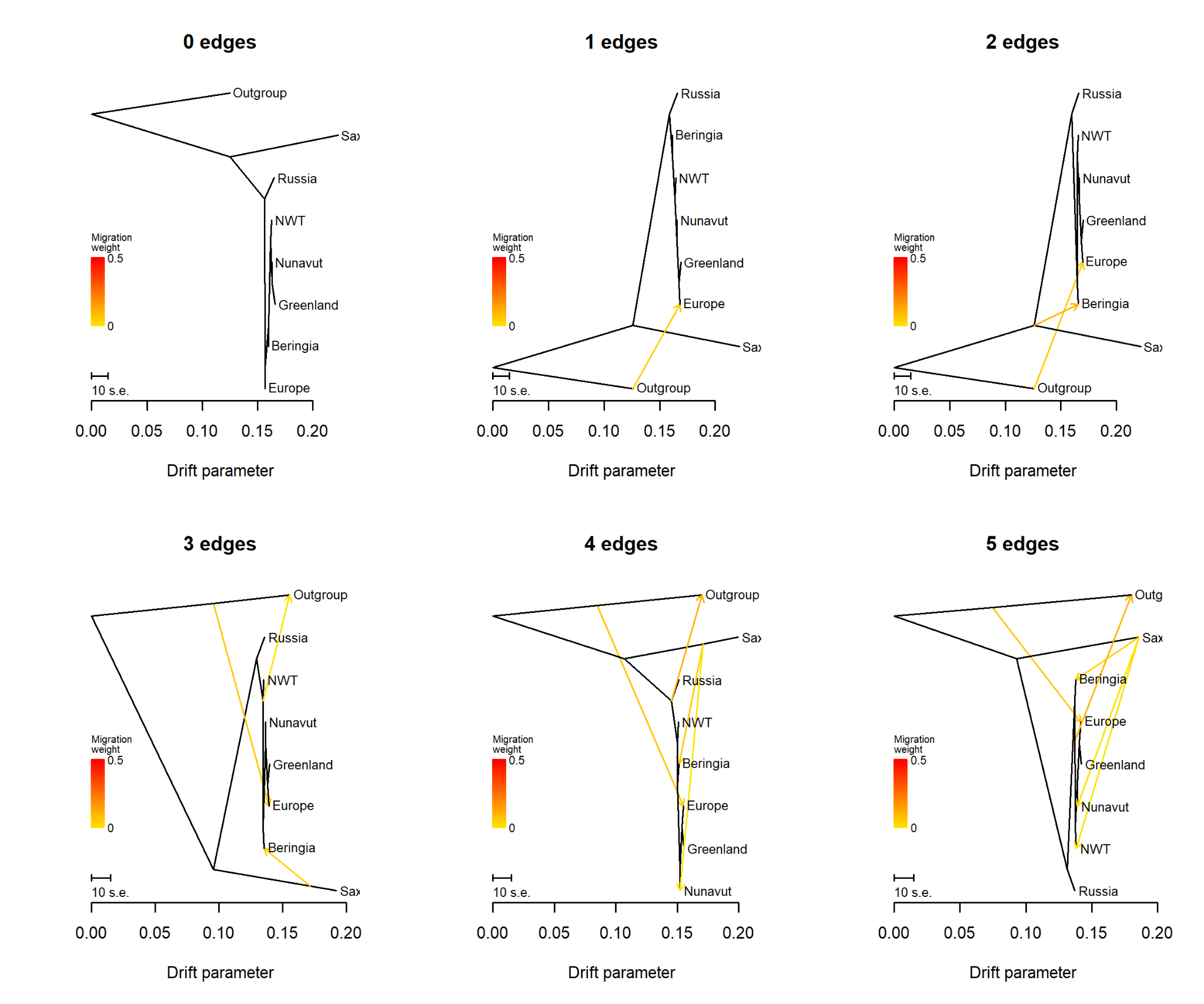

##### **Figure S10:** Maximum likelihood trees made using 349 individuals and 2310 SNPs in TreeMix 1.3.0 with all edges shown. Populations were grouped by their dominant cluster in ADMIXTURE K=6.

### Demographic models of site frequency spectra

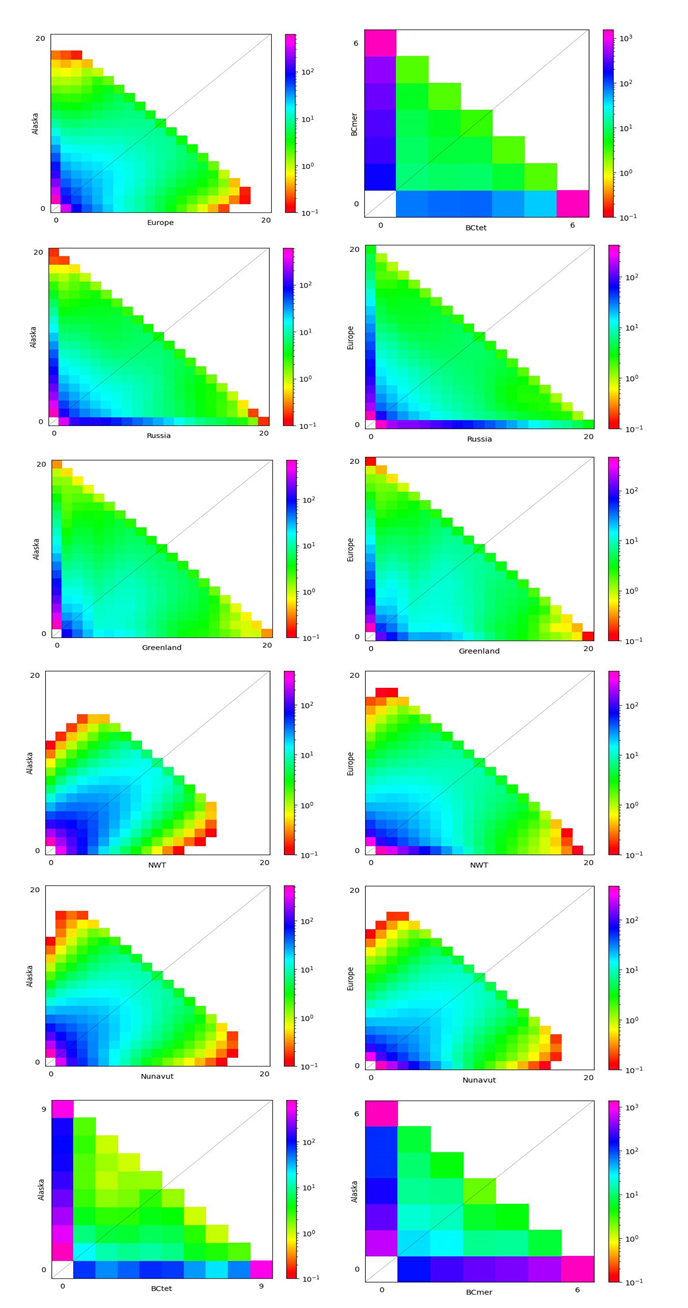

##### **Figure S11:** 2D Plots of site frequency spectra (SFS) made in dadi.

### Flow cytometry results

##### **Table S9:** Flow cytometry results for Disko Island, Greenland populations. Haploid genome size is 1.6Gbp. Both Disko populations appear to be approximately the same size.

| **Sample** | **Standard** | **Standard 2C in pg** | **Avg sample position** | **Avg standard position** | **Sample 2C in pg** | **Sample (2C) in bp** | **Sample in 2C Mbp** | **Sample 2C in Gbp** | **Sample 1C in Gbp** |
| --- | --- | --- | --- | --- | --- | --- | --- | --- | --- |
| Cassiope - DQG | Tomato | 1.96 | 52 | 30 | 3.40 | 3322592000 | 3323 | 3.32 | 1.66 |
| Cassiope - DLG | Tomato | 1.96 | 51 | 30 | 3.33 | 3258696000 | 3259 | 3.26 | 1.63 |

### AMS Carbon dating results

*C. tetragona* samples were sent for Accelerator Mass Spectrometry (AMS) carbon dating. The results can be found here:
<https://drive.google.com/file/d/1oIR-7Zwb9VJOU_ZrS98hPnfHp29Q6FHC/view?usp=sharing>

##### **Table S10:** Radiocarbon results for *C. tetragona* samples collected at Alexandra Fiord and Sverdrup Pass, Ellesmere Island, Nunavut. Calibration was performed using OxCal v4.3 (Bronk Ramsey, 2009) and the IntCal13 calibration curve (Reimer et al., 2013). Material codes are described in Crann et al. (2017).

| **Year Glacier Melted** | **Collection site** | **Material** | **14C yr BP** | **(+/-)** | **F C14** | **(+/-)** | **Range of calibrated years before present** |
| --- | --- | --- | --- | --- | --- | --- | --- |
| 2017 | Alexandra Fiord (AF) – SW Glacier | wood | 280 | 18 | 0.9657 | 0.0021 | 429-377(46.4%) |
|  |  |  |  |  |  |  | 321-290(49.0%) |
| 2017 | AF – Glenda’s Foreland (GF) | wood | 155 | 19 | 0.9809 | 0.0023 | 284-1(95.4%)* |
| 2017 | AF – Middle Foreland (MF) | wood | 150 | 19 | 0.9815 | 0.0023 | 283-5(95.4%)* |
| 2017 | AF – Adam’s Foreland (AF) | wood | 121 | 26 | 0.9851 | 0.0032 | 271-11(95.4%)* |
| 2012 | AF – Glenda’s Foreland (GF) | wood | 123 | 19 | 0.9847 | 0.0024 | 270-12(95.4%)* |
| 1992 | AF – Glenda’s Foreland (GF) | wood | 132 | 18 | 0.9837 | 0.0022 | 271-11(95.4%)* |
| 1980 | AF – Glenda’s Foreland (GF) | wood | 147 | 18 | 0.9819 | 0.0022 | 283-5(95.4%)* |
| 1959 | AF – Glenda’s Foreland (GF) | wood | 110 | 19 | 0.9863 | 0.0023 | 266-22(95.4%)* |
| 2004AF(AF) | AF – Adam’s Foreland (AF) | wood | 121 | 19 | 0.9851 | 0.0024 | 269-14(95.4%)* |
| 2017SP | Sverdrup Pass | wood | 436 | 19 | 0.9472 | 0.0023 | 521-480(95.4%) |
| *Seuss Period |  |  |  |  |  |  |  |

Carbon dating from past studies indicated that ancient samples collected in 1981-83 were 400+/- 140, 410+/-45, and 430 +/- 90 radiocarbon years old (Bergsma et al., 1984). In 1994, after the glacier had retreated roughly another 120 m, they were 830 +/- 35 years at 30 m from the 1994 glacial base, 240+/- 65 years at 70 m from the glacier, and 260 +/- 65 years at 240 m from the glacier (Jones, 1997). It is likely that the older dating from the current analysis is correct and that the plants are between 430-520 years old based on the dates when glaciers expanded on Ellesmere Island.

### ABBA BABA test

An ABBA BABA test was run in Dsuite to determine the amount of recent gene flow into the Alexandra Fiord population. The resulting D-statistic compares the amount of an ABBA configuration to that of the BABA configuration in four ordered populations, P1 through P4. To understand the two configurations, imagine P1 and P2 share a most recent common ancestor, P3 is more distantly related to P1 and P2 and P4 is the ancestral outgroup. If there is no gene flow between P3 and P1 or P2, we expect P1 and P2 to share some derived alleles not found in P3. This would appear as BBAA (P1,P2,P3,P4) where B is the derived allele and A is the ancestral allele. If there is gene flow between P1 and P3, we could get BABA and if there is instead gene flow between P2 and P3, we will get ABBA.

We also estimated an ABBA BABA f statistic which describes the proportion of genome shared due to admixture. These tests were run in a few different configurations: the outgroup (ancestral group) was set as the true outgroup *C. mertensiana,* P1 was chosen to be the ancient population from Alexandra Fiord, P2 the present-day population from Alexandra Fiord, and P3 varied with populations from Greenland, Alaska, and Europe. The ABBA configuration would occur when P2 (present day Alexandra Fiord) and P3 (Greenland) share SNPs not found in both the ancestral group and P1 (the ancient plants). The BABA configuration results when P1 (the ancient plants) and P3 (Greenland) share derived SNPs not found in the present-day Alexandra Fiord population and the outgroup.

##### **Table S11:** ABBA BABA results for each potential gene flow scenario with an outgroup (ancestral group) set as *C. mertensiana*. P1, P2, P3, refer to population placements in Figure D-1. Test results for different population scenarios are shown.

<https://docs.google.com/spreadsheets/d/1SqkSMOfKzwpvAxs0X5yc78JY8Tw7Ox7ObgrvzwmNX7M/edit#gid=39993040>

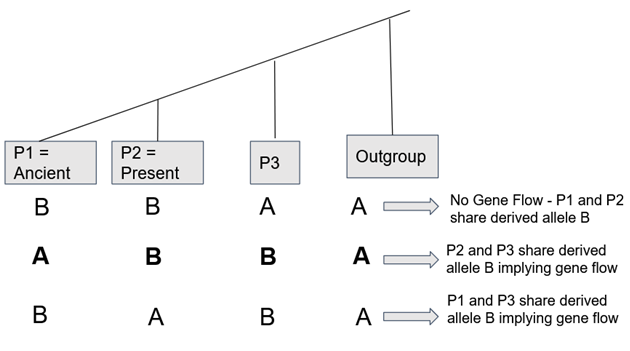

##### **Figure S12:** ABBA BABA tree where ABBA configuration (2^nd^ row, bold) supports gene flow between P3 and present-day population (results see Table D-1).
